# RegenPGC View: Semantic Segmentation of Perennial Groundcover Cropping Systems to Restore American Croplands

**DOI:** 10.1101/2025.07.03.663038

**Authors:** Bryce Meyering, Brandon Schlautman

## Abstract

Modern, conventional row crop agricultural production relies on clean tillage of croplands and bare soil during the dormant season. While this paradigm of crop production has undoubtedly led to great increases in grain yields and efficiency, it has also resulted in significant soil erosion, groundwater contamination, degradation of local ecology, and hypoxic deadzones in US watersheds. Cover cropping with perennial plant species has been proposed as a way to mitigate these negative effects of crop production while having a minimum impact on crop yields. Measuring establishment of these perennial groundcovers (PGC) in research trials is subjective, tedious, and time-consuming when calculated with traditional methods whereas image based analyses are objective, efficient, and reproducible. For this project we have developed a deep learning approach using state of the art CNN architectures to estimate PGC establishment in research plots using a variety of open-source and internal image datasets. Our novel approach uses region of interest (ROI) markers in the field, to bound the predictions which improves upon other methods. We deployed the models on AWS Sagemaker serverless endpoints, and built a lightweight Django web application to host the images and inference services. Researchers will be able to acquire plot images with smartphone cameras and get fast, reliable data from their research trials using this “Local Sensing” data collection approach. We envision that this framework can be used by other researchers and growers as PGC adoption spreads throughout the Midwestern crop production areas.

## 1. Introduction

Agricultural production has been and remains one of the defining elements that make up the bedrock of modern civilization [13]. It is undisputed that advancements in fertilizer, agricultural implements, and widespread used of agrochemicals have led to drastic increases in crop productivity [61], land use efficiency [49], and have fueled the population boom across modernizing and developing countries in the 20th century post WWII [19]. These patterns have undoubtedly led to freeing up of human labor - in the United States alone, between 1910 and 2000, the percentage of the population working in agriculture drastically declined from 14% to 1%, even as the total population increased from 92 million to 281 million people [63]. Less land is now required to sustain humanity on a *per capita* basis than at any other time in history[48].

However, increases in productivity in the 20th century have had far reaching consequences that will affect the world population throughout the 21st century if not addressed [44]. Modern practices of clean tillage and bare soil have caused increased rates soil erosion on a global scale, directly threatening food production in some of the most productive regions of the world [45, 25, 26]. Agricultural runoff containing agrochemicals, excess nutrients, and soil particles contributes to hypoxic “dead zones” in major watersheds around the globe. In the US, the Mississippi River Valley is especially impacted due to its location in central production regions prone to severe soil erosion. This runoff contributes to eutrophication in the Gulf of Mexico, which now has the second-largest dead zone in the world [46].

While early 20th century researchers boldly claimed that soil is “…the one resource that cannot be exhausted; that cannot be used up.”^1^, as early as 1943, contrarian researchers started noticing that soil erosion was an emerging problem which was drastically reduced in agricultural land that remained covered with vegetation and crop residue [20]. Throughout the middle of the century, experiments in winter cover cropping reported positive benefits on soil structure and reductions in erosion while maintaining or only slightly decreasing overall grain yields [6, 39, 9, 37]. The benefits to annual cover cropping are evident, but barriers such as increased maintenance, financial burdens associated with annual seed costs, along with the risk of implementation and lack of supporting technological infrastructure have prevented widespread adoption to these systems [2, 17].

Perennial cover crop species, here called “perennial groundcovers” (PGC), are low-growing, multi-year cover crops that can be incorporated into crop production systems. While PGC have been used historically in other perennial cropping systems such as fruit orchards [59, 62], they have not been used extensively in annual grain cropping systems. PGC have added benefits over annual cover crops since they are planted once, and maintain continuous cover on the land throughout the entire calendar year [51]. PGC have been demonstrated to have minimal impacts on corn yields and positive effects on soil water use efficiency by the cash crops [66, 67]. Furthermore they have demonstrated uses in corn bioenergy production systems mitigating the negative effects of corn stover removal [4, 3].

As with any emerging technology, there are methodological questions that arise. The RegenPGC group, a USDA/NIFA funded initiative was started to develop integrated PGC systems for annual row cropping agriculture and answer some of these questions by investigating the main barriers to widespread adoption, and aid in de-risking the systems we develop^2^. Key to the success of PGC are establishment factors such as seeding depth, emergence competition with the cash crop [28] and ensuring that PGC/corn systems don’t induce negative shade avoidance responses [51, 40]. Chemical suppression, when applied at the correct timing, can temporarily reduce corn/PGC competition and is actively being investigated [5]. Additionally, certain species under evaluation have summer dormancy, and exhibit active growth only after the corn has been harvested and thus naturally do not compete with cash crops during the growing season [65, 11]. Formulating the best management practices that allow farmers to plant, establish, and properly manage PGC systems while mitigating any potential negative impacts is a key goal within the RegenPGC project.

Getting accurate estimates of PGC growth, weeds, and bare soil coverage throughout the year are crucial KPI data for this emerging system. However these are difficult to collect, and rely on highly subjective rating scales which vary from person to person. Moreover, more accurate, quantitative methods rely on tedious measurements using sampling quadrats and intercept lines. Traditional computer vision applications based on morphological operations or simple machine learning models have been used to estimate vegetative canopies in a wide variety of crops [24, 27, 7] and have led to the development of smartphone applications such as Canopeo [42]. But these methods are coarse tools limited to identifying vegetation, generally, and cannot be restricted to certain crop classes. Recent advancements in machine learning provide a unique opportunity for technical leaders in the agricultural sector to leverage deep learning model architectures such as EfficientDet [58] and DeepLabV3+ [12] to accurately detect and segment objects in agricultural images with great success.

This current project seeks to develop two deep learning models to detect the boundaries of PGC plots, and then segment and predict important classes of vegetation in PGC field based images taken on a smartphone in “local sensing” approach. These predictions are normalized to a standard ROI and are thus directly comparable across field plots in a trial and across trials. Furthermore, a goal of this project is to develop an end-to-end image analysis pipeline, here called “RegenPGC View” that can take in a series of images, process them, and output the vegetation estimate results in a format accessible to agricultural researchers and farmers. Finally, we propose to develop an easy to use web application that ties the image management, pipeline, and data curation in one place. We anticipate that these tools will be an asset to the PGC research community, allowing them to get fast, objective data from their research trials, and informing policy and decisions about PGC systems.

## 2. Methods and Problem Approach

### 2.1. Data Sources

Several open source datasets were employed in this project. The CropAndWeed dataset (CAW) [55] consists of field-based, overhead images of 74 different crop and broadleaf weed species with a variety of different soil and lighting conditions. This dataset contained 8034 images collected over a period of 4 years in agricultural production fields in Austria. All images are fully labeled with binary masks. The full dataset is available on [GitHub](https://github.com/cropandweed/cropandweed-dataset) and can be downloaded using a Python script.

The GrassClover dataset (GC) [53] published in 2019, contains a mixture of developed and synthetic images. The developed imageset consists of ≈ 31k images taken with 3 different camera setups in Denmark. The developed data is unlabeled but consists of a wide variety of grass species, clover species, and broadleaf weeds. The synthetic dataset consisted of 8000 images that were generated from the developed images. Each synthetic image was mapped with a binary mask corresponding to the plant categories. The the developed and synthetic images can be downloaded in a zipped file from the project [website](https://vision.eng.au.dk/grass-clover-dataset/). Additionally, we used 1035 images of Kura clover (*Trifolium ambiguum* L.) breeding lines from our breeding program at the Land Institute acquired during the summer of 2017. Each image consisted of a single clover plant with a PVC sampling quadrat, and every image includes a labeled binary mask for the classes ‘soil’, ‘quadrat’, and ‘clover’. The corners of the PVC quadrats are labeled with bounding boxes.

Our target dataset, dubbed the ‘RegenPGC Dataset’ consisted of 4,192 images collected in field trial plots in corn production systems over the growing seasons of 2022-2024. The images were taken overhead, focusing on each experimental plot at approximately a height of 4 feet from the ground using a Google Pixel 3A smartphone. Some images were taken in field trials with current corn production whereas others were acquired in PGC establishment trials without any cash crop present in the field. The vast majority of these plots also have a defined region of interest (ROI) which is demarcated by the presence of four, 6” landscape fabric stakes spray-painted electric blue, or another bright color, and placed at the corners of the desired ROI. Having a permanent ROI in place for each plot allows the researchers to image the exact same portion of ground throughout the growing season in order to reduce the noise in the data. Because the placement of permanent markers was not feasible for all field trial locations considering the types and size of farm equipment available, some researchers opted to use a PVC sampling quadrat placed on top of the PGC between the corn rows to acquire images. For the marker detection, bounding boxes were drawn around the perimeter of each colored marker and PVC quadrat corner.

### 2.2. Development Environments

All image labeling was conducted using the Labelbox data annotation [platform](https://app.labelbox.com/home). Images were stored in an Amazon AWS S3 bucket for easy integration with other platforms. Image preprocessing work was conducted in Python 3.11.8 [47], using a mixture of OpenCV [8], scikit-image [64] and scikit-learn [43]. Deep learning models and training pipelines were developed in PyTorch 2 [41] and made use of the Python modules [effdet](https://github.com/rwightman/efficientdet-pytorch) [68] and [segmentation models pytorch](https://github.com/qubvel/segmentation_models.pytorch) [30] for the EfficientDet and DeepLabV3+ prebuilt-model definitions, respectively. All development work was conducted in an Ubuntu Linux 22.04 environment. Model training took place in a variety of environments. Preliminary testing was conducted in Python 3.11.8, on an Nvidia RTX 4060 graphics processing unit. Baseline segmentation models were trained on an Nvidia A100 GPU in a LambdaLabs hosted cloud server [31]. Full segmentation models were trained on an Nvidia H100 GPU also hosted in a LambdaLabs cloud server. The EfficientDet model was trained on an A100 notebook instance in Google Colab [23]. Models were evaluated using appropriate metrics from the `torchmetrics` library [16].

### 2.3. Image Labeling and Preprocessing

While the GC and CAW dataset were prelabeled by their respective authors, the Land Institute Kura clover dataset and the RegenPGC dataset required marker/quadrat labeling as well as segmentation masking. These were labeled in the online software, Labelbox. The Kura clover images (1035 images) were quick to label due to the low complexity of the images (i.e. one plant per image, bare soil, and a single white, PVC sampling quadrat). Furthermore Labelbox has an auto-detect masking feature which runs the Segment Anything (SAM) zero-shot segmentation model [29] from bounding box or point prompts, and greatly speeds up labeling projects. However, images in the RegenPGC dataset are much more visually complex due to the presence of broadleaf weeds, PGC grass and clover, corn plants, and crop residue, with ambiguous boundaries objects of different categories. Using SAM on these image features was more difficult and required careful prompting to generate the desired masks. Due to these reasons, only 282 representative images were fully labeled in the software and the remainder were left unlabeled. These images were sampled randomly from the dataset using groups output by the UMAP (Uniform Manifold Approximation and Projection) clustering functions in Labelbox.

A total of eight categories were selected for the segmentation masks: ‘background’, ‘quadrat’, ‘pgc grass’, ‘pgc clover’, ‘broadleaf weed’, ‘maize’, ‘soybean’, and ‘other vegetation’. The GC and CAW datasets each contained labels that could be aggregated up to one of these classes, so all four datasets were mapped to a common set of categories that could be applied to any image without any loss of necessary distinction. While the category ‘other vegetation’ is not a target category of interest, the CAW dataset included non-weed target crops such as sunflowers, sugarbeets, and peas that will not be present in the target dataset but should not be modeled as weeds. Crop residue (remnants of corn and soybean stubble left in the field from the previous crop season) is potentially an important category since its presence in the field can reduce soil erosion from rain events. However, since crop residue was present in the CAW dataset but was not masked as its own category, it was excluded from the list of classes as relabeling the residue in all 8034 CAW images was not feasible during the timeframe alloted to this project.

The unified, remapped images and masks were split into two main training categories: labeled and unlabeled. The total number of labeled images was 17,451 and the unlabeled portion, which included the GC developed images and the unlabeled RegenPGC images, totaled 35,844 images. The total number of pixels in the labeled dataset belonging to each category was tallied in addition to the number of images containing a given class, regardless of the number of class pixels in that image. At the same time the RGB channel means and standard deviations were calculated using the Welford method, an effective, online algorithm that avoids numerical errors due to machine *ϵ* in large sample sizes [18]. The distribution of classified pixels was heavily imbalanced across the label space (figure 1) with the top 4 categories accounting for almost 99% of all pixels in the labeled data.

**Figure 1:**
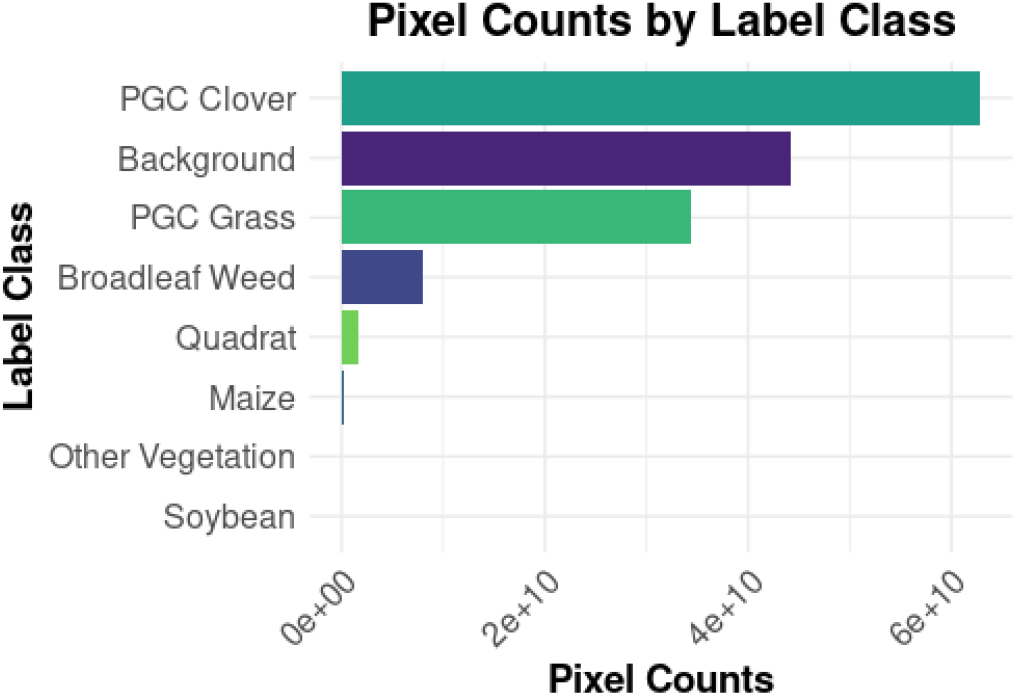
Class distribution across all pixels for the labeled dataset

**Figure 2:**
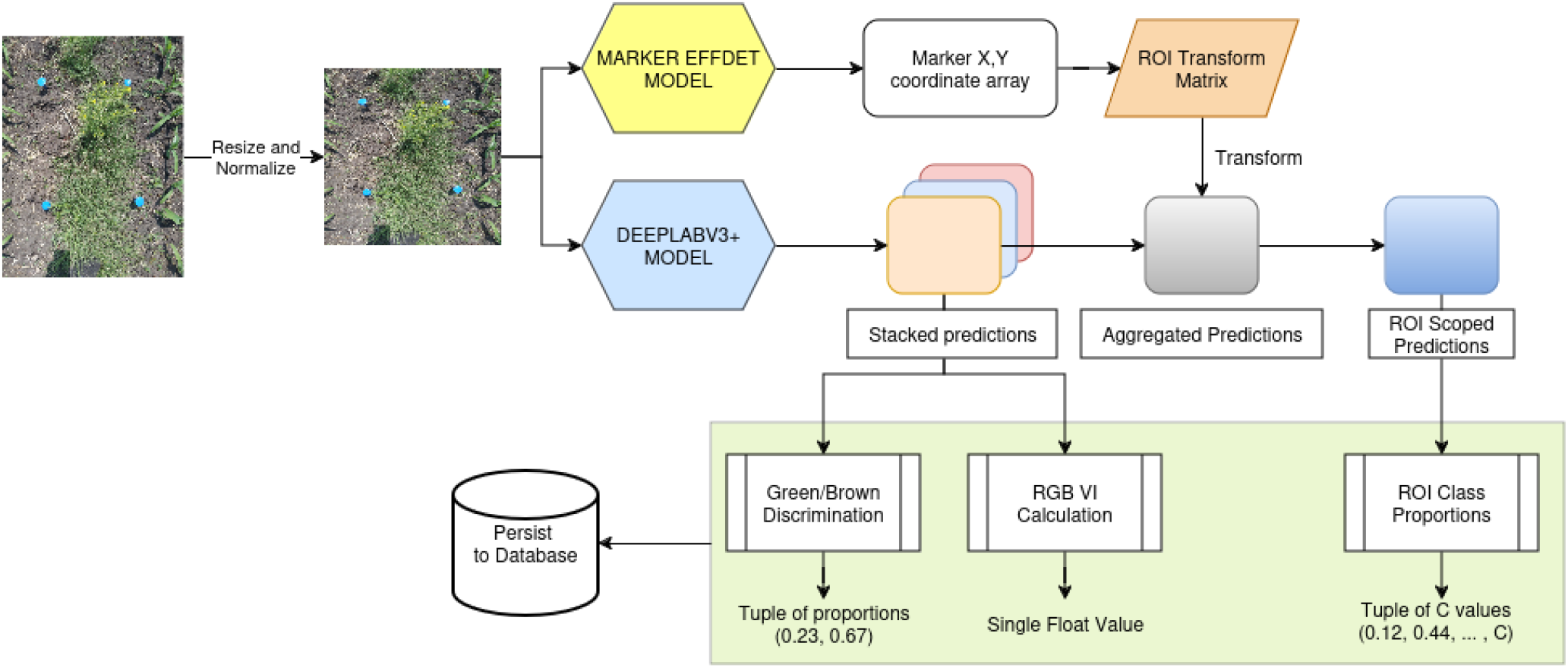
Representation of the major steps in the RegenPGC View image analysis pipeline.

The labeled images were split into train, validation, and test splits by clustering the images on the class pixel counts per image using HDBSCAN, and then stratifying the splits along the assigned clusters. This resulted in almost identical pixel class distributions in each split. The final split counts were roughly 10, 18, and 72% for the test, validation, and training splits, respectively. Additional preprocessing such as normalization, resizing, augmentation, were performed on the images during modeling but were only performed on the fly and not saved to the images.

### 2.4. Marker Detection Modeling

Detection of the plot ROI corners, whether determined by the colored landscape markers, or the PVC quadrat corners was carried out by training an EfficientDet single shot detection (SSD) model in Pytorch. The EfficientDet model uses a combination of multi-scale feature fusion and model scaling over multiple dimensions using a bi-directional feature pyramid network [58]. These innovations combined with more modern backbones such as the EfficientNet [57], allow for greater model accuracy with far fewer FLOPS than other SSD model architectures. Our marker detection model was trained with an EfficientNet D5 backbone with an image input size of 1024×1024. We modeled all of the round colored markers as the same object class and added in the quadrat corners as an additional class. The network training was optimized with an ADAMW optimizer [34], and an initial learning rate of 0.005 with an exponential learning rate scheduler. The model was set up to train for a total of 60 epochs but used an early stopping callback function in Pytorch Lightning with a patience period of 5 epochs. The effdet models trained were evaluated using average precision (AP) and average recall (AR) at different scales on the validation datasets.

### 2.5. PGC Segmentation Modeling

Semantic segmentation modeling can prove to be a difficult task, especially in situations with extreme class imbalance, such as what our labeled dataset exhibits. We broke this problem down into several sub-problems that allowed us to pinpoint the best solution to the problem more effectively. 1) Train a series of preliminary models using 11 different loss functions aimed to address the class imbalance problem. 2) Train better baseline models using the top *k* loss functions but using a better backbone and a higher image resolution. 3) Implement the FixMatch [54] semi-supervised learning (SSL) algorithm to incorporate the most confident predicted pixels of the unlabeled data to iteratively improve the performance of the model.

#### 2.5.1 Loss Function Implementation

We implemented a series of 11 different loss functions to choose the most performant ones given the class imbalance in the dataset. As a rule, we always included a vanilla (unweighted) cross entropy (CE) loss in all of our comparisons. The Python library `segmentation models pytorch` comes preloaded with several of these loss functions, but it has a largely inconsistent API and doesn’t have a pixel masking feature for all of the losses, which is crucial to the FixMatch algorithm. So we took some insights from their design and implemented them in this project from scratch, except for vanilla CE loss for which we wrote a wrapper around the official PyTorch implementation in order to conform to our call design. We chose a mean reduction strategy across all pixels and images in a given batch.

##### Vanilla CE

One of the most common loss functions for classification, multiclass cross entropy, CE is defined as

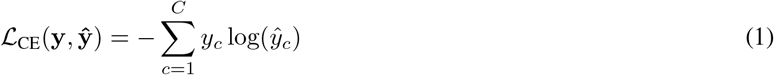

where *y*_*c*_ is the one hot encoded vector of ground truth class assignment for a given pixel and log *ŷ_c_* is the vector of *C* softmax probabilities for that pixel which sum to 1.

##### Inverse Weighted CE

Inverse weighted CE loss takes the above definition a bit further and allows us to weight each class differently based on any weight criterion that we define. WCE is defined as

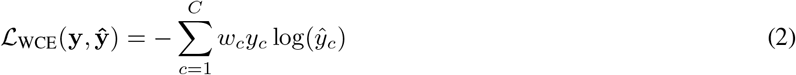

where we introduce a class weight term *w*_*c*_ for each class *C*. For our implementation, the weights can be passed directly to the class upon instantiation. The inverse weighting strategy uses the total number of ground truth pixels for a given class *N*_*c*_ in the training data which is shown in figure **??**. These are are all divided by the sum, *N*, of the number of pixels in all classes. Finally, we take the inverse and arrive at 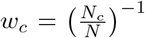. This forces the model to weight mistakes in the minority classes higher than those of the majority classes.

##### Focal Loss

Focal loss (FL) [32] introduces a focusing parameter *γ* which determines how aggressively to downweight samples that are easy to classify. This term is added in as an adaptive weighting term on the *1 − ŷ_c_* and is defined as

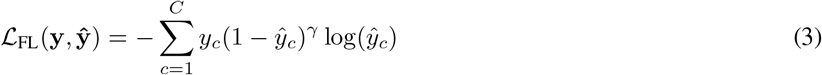

When the model is very confident for a given prediction *ŷ_c_* and *y*_*c*_ ≠ 0, i.e. it is the correct prediction, this loss value does not contribute that much to the overall loss. Typical values for *γ* are either 1 or 2. Note that FL can easily incorporate a weight term as in eq. 2 to arrive at

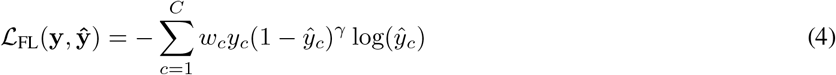

##### Class Balanced Loss

CB Loss takes the focal loss a bit further by recognizing that the inverse weighting methods tend towards type I errors since the models tend to overfit the minority class data [15]. This loss defines a new weighting term, the effective sample number 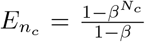 where *N*_*c*_ is the number of samples in class *C*, and *β* is a user set hyperparameter.

In the paper, they rescale each weight term such that 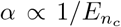 and 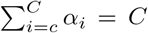 which keeps the weights in a more manageable range. CB loss is then calculated as

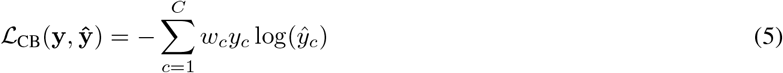

Where 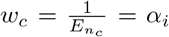. Since this definition is just an extension of eq. 2 and eq. 4, we wrote this class as a wrapper that calculates the new weighted terms and passes them to the WCE and FL loss class implementations.

##### Adaptive Class Balanced Loss

ACB loss adpated the CB loss above by instead letting *β* be dynamically calculated from the data above instead of relying on (possibly) expensive and time-consuming hpyerparameter tuning [69]. To this end, several new terms *u, v*, and *b* are defined from the dataset itself and mapped back to *β*.

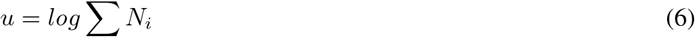

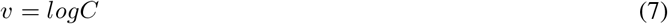

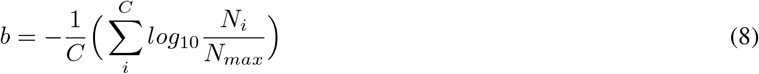

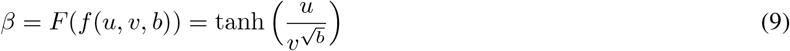

Finally 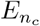 is calculated from *β* the same as in eq. 5 for CB loss and is thus available to use as a vector of *C* weights in both CE and FL.

##### Recall Loss

Tian *et al*. found an additional way to reweight cross entropy loss as Recall Loss (RL) by defining the recall weights as the false negative rate for class c [60].

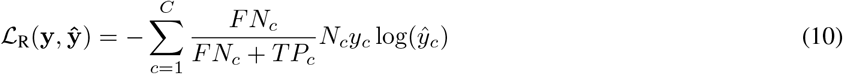

And seeing that Recall = 1 - FNR, the, Recall CE loss, for step *t* in training is calculated as

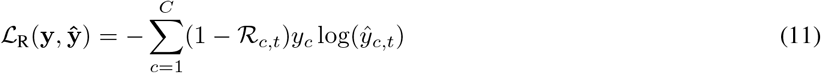

thus the loss weights are dynamically changed to the reflect the most recent performance of the model at time *t*_*n*_ instead of keeping static weights that are created from the label distributions at *t*_0_.

##### Dice Loss

Segmentation models generally measure model performance using either the Dice Score or the Jaccard Index (IOU) which measure the amount of overlap between the ground truth and the prediction. However, these metrics are non-differentiable and thus cannot be used in network training during backpropagation. The Dice Score can be defined as follows for two object areas *A* and *B*

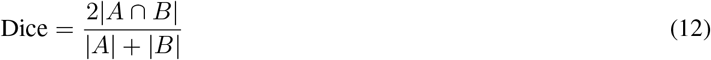

Which in terms of the predictions of the pixel classes can be equivalently written as

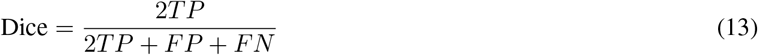

Being able to directly optimize the model for the score used to evaluate the model is a judicious insight, leading researchers to develop a differentiable version adaptable to multiple classes [56].

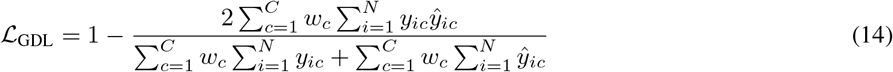

Which can easily accept new weights *w*_*c*_ and is suitable for direct usage in optimization during backpropagation.

##### Tversky Loss

Finally, the Tversky Loss (TL), based on the Tversky similarity index, allows one to set hyperparameters *α* and *β* to weight the FP and FN differently and generalizes both the Dice Score and Jaccard Index in one differentiable loss function that is adaptable for multiclass segmentation problems [50]. The Tversky similarity score is defined as

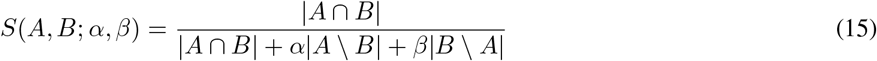

where |*A \ B*| is FP and |*B \ A*| is FN. Then the differentiable Tversky Loss can be written as

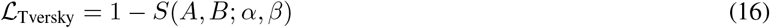

Which is expanded as a multiclass function as the following formulations:

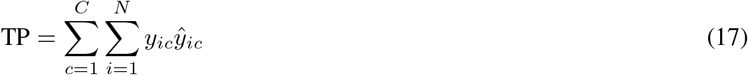

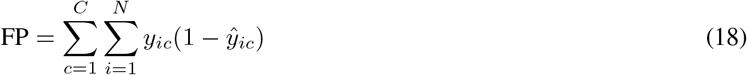

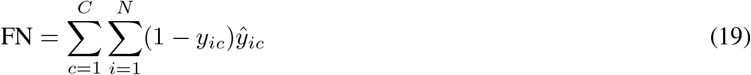

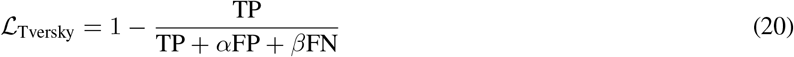

It is noted than when *α* and *β* are both set to 0.5 the TL converges to GDL, but when not equal, allows one to leverage different weights for FP and FN.

Considering that CB, ACB, RL are just alternative strategies to generate a vector *w*_*c*_ of weights for each class *C*, these can be directly incorporated into both WCE and FL. This consideration gives us room to evaluate 11 different loss functions. We designed each of these loss classes in Python with a standard, consistent argument signature, so they could easily be used interchangeably for rapid testing. Additionally, each of them accept a boolean `mask` parameter that allows one to discard any loss values from non-confident pixel predictions during FixMatch training, which was missing in several other released Python implementations of these online.

#### 2.5.2 Preliminary Model Training

Preliminary, fully supervised models using only the labeled dataset were trained and optimized with the loss functions defined above. The purpose of the preliminary training was to quickly generate performance scores across the metrics using low resolution images and a smaller backbone and select the top *k* performing loss functions. To this end we trained 11 DeepLabV3+ models using a ResNet50 encoder backbone with pretrained Imagenet starting weights. The learning rate was set at 0.001 and incorporated into an exponential LR scheduler. Optimization was performed using SGD with Nesterov momentum. An L2 regularization weight decay of 1e-5 was applied to all parameters except batch norm and biases. Additionally, our models employed an exponential moving average (EMA) of model weights as additional regularization with an EMA decay of 0.9 to reduce the noise in the weight updates [38]. The image size was reduced from the raw resolutions, typically in the range of 3000-4000 px width, to a much smaller 512×512. A standard set of image augmentations including randomized rotation, flips, affine shifts, and Gaussian blurring were applied on the fly during training, and each image was normalized to have mean 0 an *σ* = 1 using the RGB means and *σ* calculated on the training data. Each model trained for a total of 10 epochs, and the models were evaluated during training and validation using the F1-score and Jaccard Index, both as the global average over all classes, as well as for each individual class.

#### 2.5.3 Baseline Model Development

A total of four baseline models were trained using the same fully labeled dataset and loss functions from the top three models including vanilla CE model that was included as a non-class weighed comparison model. We kept the DeepLabV3+ model decoder but switched to a more modern ‘efficientnet b4’ backbone. The backbone was initialized with pretrained imagenet weights. The image size was increased from 512×512 to 1024×1024, and each model was trained for a total of 20 epochs using the same LR, EMA, L2 regularization parameters, and image augmentations from the preliminary model training. Checkpoints were saved only in the event of a decrease in total validation loss. Model evaluation metrics were kept the same and were logged to Tensorboard after each epoch. Loss curves were logged at both the batch step as well as the epoch average. The best checkpoint for each model was used as the set of starting weights for the FixMatch implementations.

#### 2.5.4 FixMatch Implementation and Training

We implemented a custom FixMatch training class in order to evaluate the effectiveness of using the learned model parameters from the fully labeled datasets on unlabeled data of unknown quality which we based upon the published implementaiton [54]. In brief the FixMatch algorithm includes two sources of loss that get combined into one total loss value for network backpropagation. The first loss term, ‘labeled loss’, is the direct, unchanged output from a given loss function when a fully labeled batch of images runs through the network during the forward pass. The second term, ‘unlabeled loss’, is more complex, but yields an intuitive result. First, a batch of unlabeled images goes through a series of weak augmentations including randomized affine shift, weak Gaussian blurring, and horizontal flips. The resulting transformed images are then passed through much stronger augmentations that add safe rotation, solarization, RGB channel shuffle, and RGB shifts. These two sets of images entitled ‘weak images’ and ‘strong images’ are then concatenated and passed together through the network to obtain the raw logit scores, which are then normalized through a softmax layer and an argmax function to find the best prediction class per pixel. The ‘weak image’ pixel scores are then separated from the ‘strong image’ scores. Using the threshold parameter *τ* ∈ [0, 1] from the configuration file we constructed a boolean tensor, mask_*n*_ for each weak image *n* from the tensor of `argmax` confidence values, conf_*n*_ where pixel *i, j* takes on the value

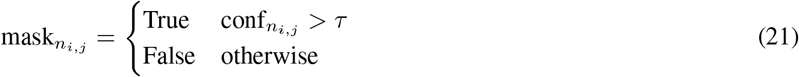

Additionally, for batch *t* we defined a weight *f*_*t*_ which is defined as the fraction of total pixels *N*_*t*_ in the batch which are above *τ*, or

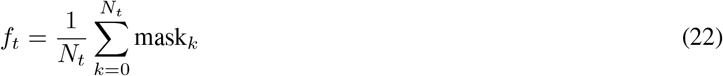

where the subscript *k* represents the position on the flattened batch mask vector.

The `argmax` predictions from the weak images are designated as the ‘pseudo-labels’ for each pixel. These pseudo-labels and the strong images are passed through the loss criterion as usual to output a loss score. Each of the loss classes excepts a boolean `mask` argument, and this gets multiplied by the pixelwise loss values, before mean reduction, to nullify any pixel whose classification confidence was *< τ*. Finally, the mean loss of the masked losses is calculated and returned as ‘unlabled loss’. In the end, the total loss value is a weighted sum of the labeled and unlabeled losses modulated by the weighting factor *λ*. The original formulation is shown below:

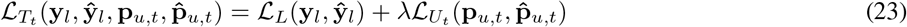

where ℒ_*L*_ and ℒ_*U*_ are the labeled and unlabeled losses respectively, and **p**_*u,t*_ and 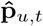 are the pseudolabels and predictions from the pseudolabels on the unlabeled dataset reflecting the capabilities of the model at time step *t*. While it is the case that the model should learn the most confident pixels from the unlabeled data, hence the masking procedure, it may be inappropriate to have a constant weight term *λ* for every batch. This is especially true in situations where you have a batch with very few confident pixels. Even though a small fraction of pixels are contributing to ℒ_*U*_, the mean reduction strategy would equally weight those few pixels in comparison to a situation where all of the pixels contain highly confident predictions.

Thus we also implemented the fractional weight *f*_*t*_ (eq. 22) which downregulates the unlabeled loss as 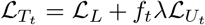. Seeing that *f*_*t*_ ∈ [0, 1] this has the effect of also handling case when there are no confident pixels in the unlabeled data and thus when the loss is calculated, it results in an `Nan` error. We can use *f*_*t*_ directly as follows:

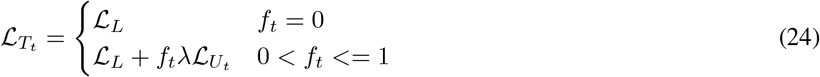

Using this method, we trained the FixMatch algorithm over the combined labeled and unlabeled portions of the dataset using the baseline model checkpoints from the previous step as a starting point. Starting from some pretrained weights helps the model converge to decent predictions quicker as starting from scratch using FixMatch can result in a lot of instability during the first epochs. These were trained for a total 25 epochs with the same parameters as the baseline models. In order to account for differences in the number of samples in the labeled and unlabeled data, we used a slightly larger batch size for the unlabeled images and also implemented an infinite sampler for the labeled data PyTorch dataloader to avoid `StopIterationError` exceptions, so there is some small but non-zero chance that each image in the labeled dataset will be sampled twice within the same epoch. Metrics were logged to Tensorboard the same as before.

### 2.6. Dormant Fraction Pixel Estimation

PGC grass dormancy changes over the season and is dependent on the grass species being trialed. We want to capture the proportion of pixels in the predicted PGC grass mask that are dormant versus the pixels that are brown and dormant at a given timepoint to allow researchers to model the phenology of PGC dormancy when grown as cover crops under corn and soybeans. This is a difficult problem to address in the segmentation model, not because it is methodologically difficult to optimize, but due to the fact that, generally speaking, dormant grass and active grass grow closely together and are extremely difficult to label accurately. Furthermore, the CAW and GC datasets include grass labels, but only green grass is represented in the images. Thus we are implementing the green/brown PGC pixel discrimination as a post-processing layer on the model outputs. We addressed this post-processing in several different ways.

First, we employed a naive approach by taking the predicted pgc grass mask and using it to filter the original RGB image for just the pgc grass pixels. We convert these pixels from the RGB colorspace to HSV space and then set upper and lower thresholds for green and brown colored pixels separately and apply them to the mask. Summing over these filtered mask pixels and then dividing by the total number of pixels in the unfiltered image mask returns the proportion of green and brown pgc grass pixels, respectively.

Secondly we approached this using a machine learning approach using Gaussian Mixture models. Flattening the HSV image masked for pgc grass we can treat each pixel as a distinct data row. We set the number of clusters to 2 and set the cluster means priors as the midpoints of HSV green and brown pixel ranges. The rationale being that most pixels in the ‘green’ range would have a higher conditional probability of being generated by the ‘green’ Gaussian process compared to the ‘brown’ process, i.e. *P* (*y* = green |**x**_*i*_) *> P* (*y* = brown | **x**_*i*_). We then set a threshold to select the green and brown pixels and generate their respective filtering masks. Since these images are inherently noisy, a side application of this would be process the masks using a simplification of the image using a superpixel generator such as SLIC [1]. This has the added benefit of smoothing regions of noise in the image and finding the distinct boundaries (if any exist) between the ‘green’ and ‘brown’ regions.

In order to facilitate both approaches, we recognized that unevenness in lighting plays a key role in image segmentation, contributing to overexposure in the lighted areas an under-exposure in the shade. This is particularly pronounced in late season images once the corn plants have formed a dense canopy over the PGC growing underneath. We attempted to mitigate some of this unevenness by converting the image to the l*a*b* color space, extracting the l channel, and applying some adaptive thresholding algorithm such as CLAHE [70]. The newly transformed channel is then reinserted into the array and converted back to RGB to generate a more lightning neutral image.

A related, but non-necessary addition to this post-processing step is the calculation of vegetative indices. Vegetative indices, or VIs, have been used extensively in remote sensing to discriminate between actively growing, healthy cropland vs dormant, or physiologically stressed cropland using satellite or UAV imagery. The best VIs incorporate NIR bands into their calculations since these healthy plants strongly reflect light in these wavelengths. When only RGB data is available we can calculate one of many different RGB VIs which can reasonably approximate NIR-RGB VIs under the right assumptions and conditions. Several papers have compiled and evaluated many of these indices so we will refer readers to those publications [33, 10, 35]. We implemented many of these in Python and use them as final return data in the image analysis pipeline.

### 2.7. Image Analysis Pipeline

The main goal of this project is to combine the outputs of the three main thrusts of this project, the marker detection model, the segmentation model, and dormant fraction pixel discrimination into one unified image analysis pipeline. This section details how this is accomplished for one image.

1. Predict plot ROI corners and calculate ROI transformation matrix.
2. Predict class masks and store as (C, H, W) binary array.
3. Discriminate green/brown grass pixel fractions and store values.
4. Aggregate prediction masks, transform, and sum the classes.
5. Persist calculations to file or database.
6. (Optional) Output intermediate images and transformations

Here we explain every step of the pipeline in detail, which is outlined graphically in 2. First, a selected image gets read in, normalized, and resized to (1024, 1024). This image is then passed through the marker detection model, either in batches, if using a GPU, or one by one on a CPU powered system. The EfficientDet marker model returns an Nx6 tensor of bounding box coordinates, their confidence, and the object classification, ranked highest to lowest in terms of decreasing confidence. The top 4 confident predictions are selected, or the top *k* predictions with a confidence above 0.5. If the model only returns 3 due to one marker being occluded or some other reason, we use some geometric methods to try to impute the position of the 4th marker. If there are fewer than 3 confident predictions, no markers can be imputed and the model will return no bounding boxes. The centers of each of these boxes are calculated and used to find the corners of the ROI. Since we know the distance of each ROI marker to the 4 corners of the image, we can calculate a 4×4 distance matrix between the image corners and the ROI corners, and then assign each ROI corner to the closest image corner using a linear sum assignment [14]. This allows us to accurately calculate a transformation matrix for the ROI using a four point transform.

Second, the resized, normalized image is passed through the DeepLabV3+ model. The output of this model is a CxHxW tensor of logits. These are passed through a softmax function such that the resulting proportions sum to 1 for each pixel along the C dimension. To obtain the class predictions, we take the `argmax` along that dimension and return a binary array of size (1024, 1024) containing up to *C* − 1 integers starting at *C* = 0 for the total number of classes in the model. The masks are stored in a Numpy array until needed later in the pipeline.

Third, convert the RGB image to l*a*b colorspace, and perform CLAHE on the l channel. Reassemble the array and convert to HSV. Extract the pgc grass prediction slice from the segmentation array and use it to filter for just grass. Run the green/brown discrimination functions to output a tuple of two float values, one for each fraction. Additionally, we calculate the RGB VIs at this point and can use the pgc grass mask to filter just the relevant portions of the image and return the average VI value for all the predicted grass in the ROI.

Fourth, take the prediction tensor from step 2, and aggregate all of the channels into one (1024, 1024) binary tensor. The index of the channel dimension, C, in the prediction tensor is mapped to the 255 values in its corresponding channel mask. So given a tensor *X* ∈ *{*0, 255*}*^*C×H×W*^, map it to a tensor *Y* ∈ [0, *C* − 1]^*H×W*^ such that: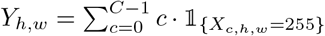. This binary mask will get transformed into the ‘top down’ view of the plot and cropped to the ROI using the ROI transform matrix from step 1. Now, the pixel classifications are limited to what appears within the ROI bounds. We can just simply tally the number of pixels in each class and normalize by the total number of pixels to get the proportion of each class in the ROI. This transformation is important not only to limit the detections to the area of the plot that we are interested in measuring, but it also transforms the shear out of the image if the image was taken at a non-ideal rotation or an oblique angle to the ground. Fifth, all of these values, the proportion of ROI area predicted for each class, as well as the fraction of green/active pgc grass and brown/dormant pgc grass and (optionally) the plot ROI mean VI value for the predicted pgc grass will then be persisted to a file or to a database depending on whether the pipeline is run locally or in the cloud. The data row gets stored with a timestamp, the image name, and other associated metadata that is important for downstream analysis.

Finally, this pipeline presents the option to export all of the intermediate images in the process. For example one might want to store the original class predictions over the entire image, the class predictions for the ROI, the original RGB image cropped to the ROI, etc. Using the aggregated mask, we can pseudo-color the binary image, back transform to the original image, and then overlay as a colored alpha channel on the original RGB image. While the image intermediate outputs are optional, these are useful to researchers who want to visualize various points within the pipeline.

### 2.8. Model Deployment

Both the marker detection model and the PGC segmentation model are deployed using AWS Sagemaker serverless end-points. These allow the models to spin up once the endpoint is pinged, scale up to the necessary number of instances to handle the inference load, process the results, and then scale back down to zero instances after a set timeout period. We expect that the workload traffic will be intermittent, thus serverless endpoints are good, cost-effective approaches to model deployment. We developed a lightweight web application using the Django development framework that will allow users to upload a batch of images and submit them as an inference job. The image preprocessing (normalization, resizing, etc.) will take place as a service layer within the application itself and then submitted to each endpoint through an AWS API Gateway. The models will run inference on each image and return the results. The results then get deserialized in the application, and passed on to the rest of the inference pipeline within the application. The RegenPGC View application has a dockerized PostgreSQL database running in the backend, to which the inference job, image metadata, and the processed results tied to each job will get persisted. Users will have a login and be able to interact with jobs that they submitted, and download the results from current and past jobs.

## 3. Experiments and Results

### 3.1. Marker Model Training

Using a subset of the RegenPGC images we trained an EfficientDet model to detect and locate either colored landscape fabric stakes in the ground or PVC quadrat corners. In total there were 1460 images in the training data set and 365 in the validation set, which represented a total of 2107 marker instances and 5094 quadrat corner instances. The marker model training loop was set to train for a maximum of 60 epochs with an early stopping callback. In practice, most of the model training runs to test out various hyperparameters ran for no more than 25 epochs after achieving high average precision (AP) and average recall (AR) at different scales on the validation dataset (Scores not shown for brevity). The model train and test loss were closely matched throughout training with no signs of overfitting. During training we evaluated backbone architectures ‘efficientnet v2’, ‘efficientdet d4’, ‘efficientdet d5’, and ‘efficientdet d6’. While the best model trained implemented the d4 backbone and achieved a validation loss of 0.113, almost all of the backbones tested had similar performance upon validation and converged easily. This was the most straightforward portion of the project since there was very little tuning required to train an accurate model, and the targets are for the most part, distinct and stand out from the surrounding vegetation.

Occasionally there are some false detections due to other objects such as flags or colored notebooks that were left in the field. Since these are rare events, many of these images were not labeled and the model didn’t learn to avoid localising them in the image. Care should be taken in future image acquisition sessions to avoid having these within the camera frame. The training set included many pictures of partially occluded markers thus the final model was able to not only locate both classes with a high degree of precision, but also performed well in scenarios when a given marker object was occluded by vegetation, which happens frequently in the field, especially near the end of the growing season (figure 3). While this worked well for the landscape markers, the same could not be said for when the quadrat corners were occluded. Since quadrats are generally placed on top of the sampling surface, and are thus temporary sampling ROIs not prone to being occluded by vegetation, this is an important edge case to be able to handle, especially when corn leaves hang over into the pgc row and could potentially occlude a portion of the quadrat.

**Figure 3:**
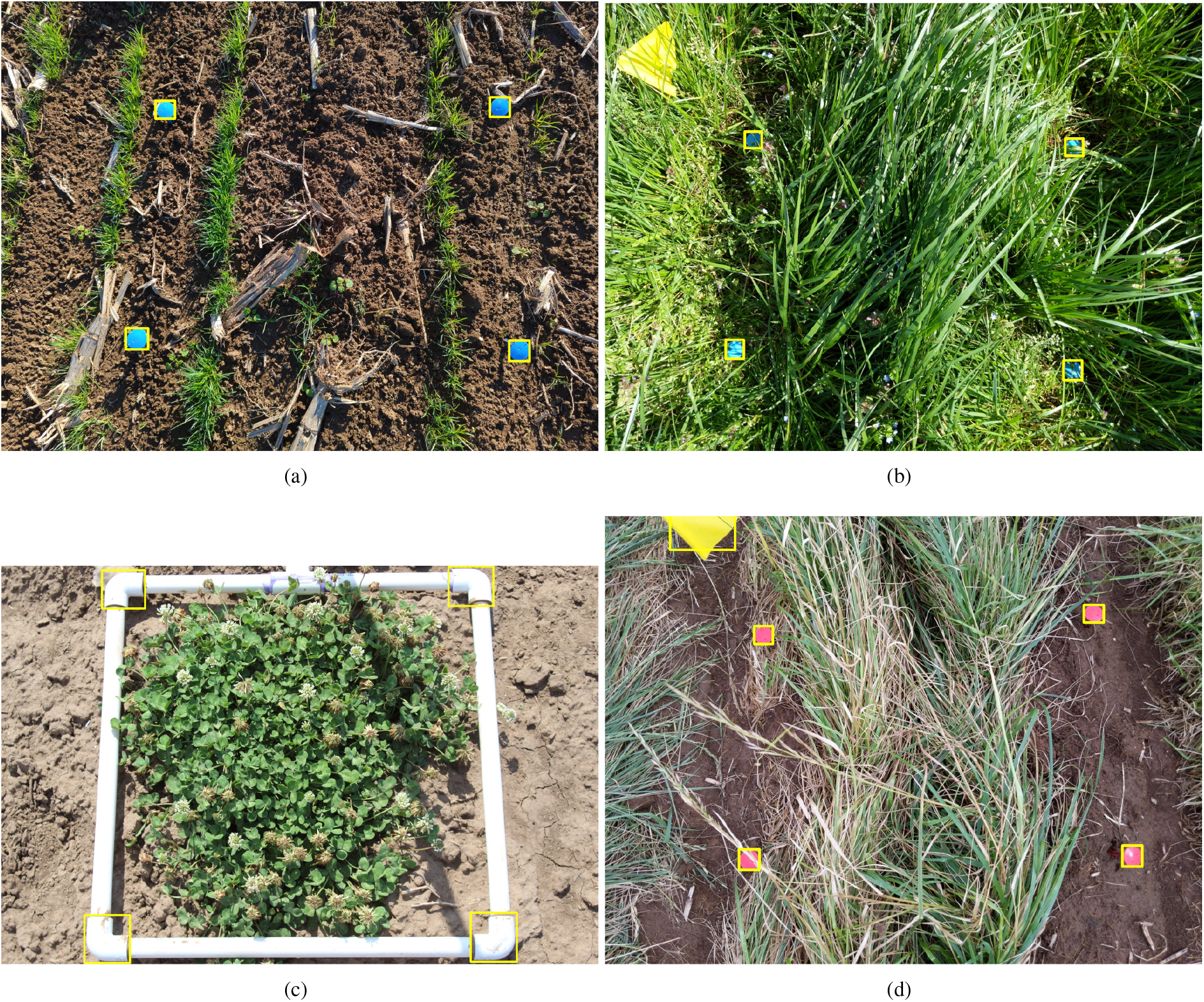
EffDet ROI marker detection examples (a) fully visible landscape markers (b) partially hidden markers (c) PVC sampling quadrat corners (d) False detection due to colored marking flag in the camera field of view

While this current model is highly accurate and more than sufficient for a version 1 of the image pipeline, it could be made more robust by injecting more difficult to localise and classify samples in different images. Regarding ROI marker classification, there was very little misclassification between quadrat corners and colored markers - the only misclassification in the validation dataset occurred due to detections with low confidence (*<<* 0.05) since the model returns the top 100 predictions. Additionally, this model was trained with markers that were blue, yellow, pink, red, and brown, however the vast majority of the marker instances in the image were blue or pink. There are opportunities to expand the model in the future to classify each marker color as a different class so that ROIs associated with certain colors could be treated differently in the pipeline. For instance, plot ROIs that contain only pgc clover do not need to be passed on to active/dormant pixel discrimination and could be handled using this feature. At the moment, we don’t have enough labeled training data for each of the separate colors to be able to handle this and thus we classified them as one category. These minor adjustments to handle edge misclassification cases, partially occluded quadrat corners, and expand the number of marker classes to accommodate unique marker colors that correspond to ROIs with different purposes are all improvements that can be expanded upon for the future after more data is gathered.

### 3.2. Preliminary Modeling for Loss Selection

Using only the labeled portion of the semantic segmentation dataset, we trained a series of 11 preliminary models with a reduced image size of 512×512 and a ResNet50 encoder backbone within a DeepLabV3+ architecture. Each model was trained for a total of 10 epochs and were generally converged well by the end of training. The best set of weights out of all epochs were selected and used to calculate the final evaluation metrics both on a classwise basis and over all classes. All of the models had converged by 9 or 10 epochs, some had converged before. Vanilla cross entropy employs no rebalancing between different classes and was used a control group to compare the rest of the loss functions. In general, CE loss performed significantly better than most of the loss functions that were supposed to help balance out the classes. CE predicted the ‘soil/background’ class well, with high overlap between the ground truth, as well as ‘quadrat’, and ‘pgc clover’. However this model did not predict anything in the ‘soybean’ class and had poor predictions in the ‘pgc grass’ and ‘broadleaf weed’. Even still CE loss produced one of the top 4 models when looking at the overall Jaccard Index (table 1) and had a reasonably high F1 score of 0.8799 (table 2).

**Table 1:**
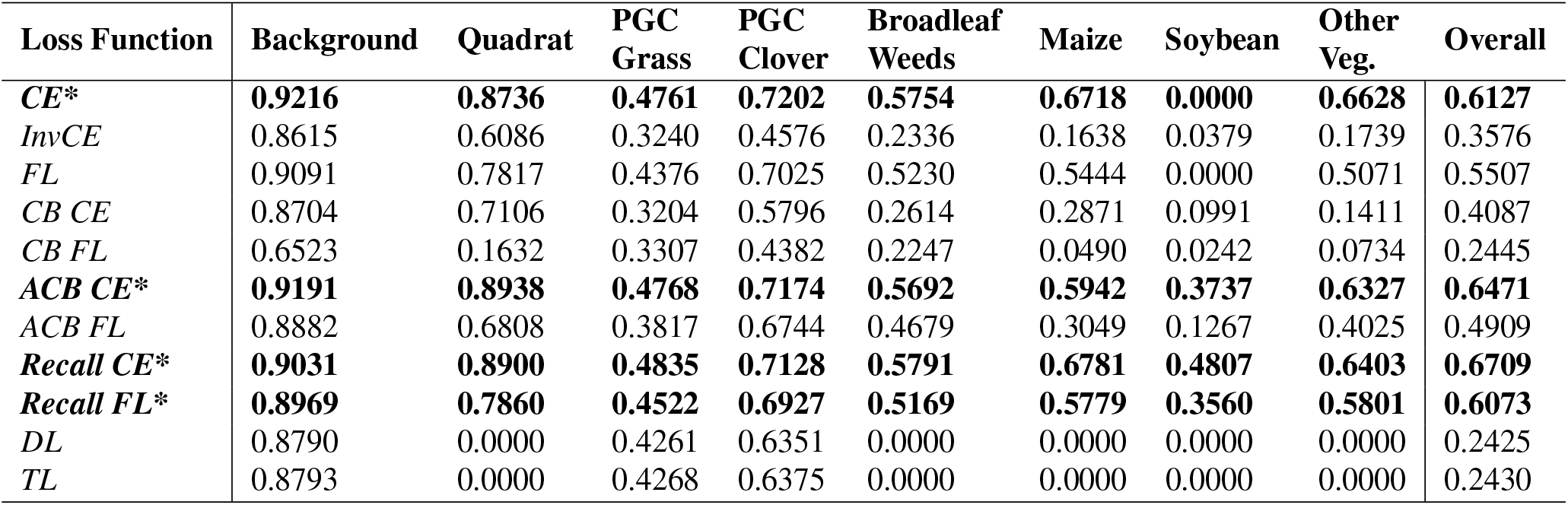
RegenPGC Segmentation Model: Classwise Jaccard Index scores for the evaluation set CE - Cross Entropy, InvCE - Inverse weighted CE, FL - Focal Loss, CB - Class Balanced Loss, ACB - Adaptive Class Balanced Loss, Recall - Recall Loss, DL - Dice Loss, TL - Tversky Loss. Jaccard Index evaluation scores span the interval [0, 1] with higher values indicating better overlap between the ground truth mask and the predicted masks. * next to a loss function and bold rows indicates this loss was selected for baseline model training.

| Loss Function | Background | Quadrat | PGC Grass | PGC Clover | Broadleaf Weeds | Maize | Soybean | Other Veg. | Overall |
| --- | --- | --- | --- | --- | --- | --- | --- | --- | --- |
| **CE\*** | **0.9216** | **0.8736** | **0.4761** | **0.7202** | **0.5754** | **0.6718** | **0.0000** | **0.6628** | **0.6127** |
| *InvCE* | 0.8615 | 0.6086 | 0.3240 | 0.4576 | 0.2336 | 0.1638 | 0.0379 | 0.1739 | 0.3576 |
| *FL* | 0.9091 | 0.7817 | 0.4376 | 0.7025 | 0.5230 | 0.5444 | 0.0000 | 0.5071 | 0.5507 |
| *CB CE* | 0.8704 | 0.7106 | 0.3204 | 0.5796 | 0.2614 | 0.2871 | 0.0991 | 0.1411 | 0.4087 |
| *CB FL* | 0.6523 | 0.1632 | 0.3307 | 0.4382 | 0.2247 | 0.0490 | 0.0242 | 0.0734 | 0.2445 |
| **ACB CE\*** | **0.9191** | **0.8938** | **0.4768** | **0.7174** | **0.5692** | **0.5942** | **0.3737** | **0.6327** | **0.6471** |
| *ACB FL* | 0.8882 | 0.6808 | 0.3817 | 0.6744 | 0.4679 | 0.3049 | 0.1267 | 0.4025 | 0.4909 |
| **Recall CE\*** | **0.9031** | **0.8900** | **0.4835** | **0.7128** | **0.5791** | **0.6781** | **0.4807** | **0.6403** | **0.6709** |
| **Recall FL\*** | **0.8969** | **0.7860** | **0.4522** | **0.6927** | **0.5169** | **0.5779** | **0.3560** | **0.5801** | **0.6073** |
| *DL* | 0.8790 | 0.0000 | 0.4261 | 0.6351 | 0.0000 | 0.0000 | 0.0000 | 0.0000 | 0.2425 |
| *TL* | 0.8793 | 0.0000 | 0.4268 | 0.6375 | 0.0000 | 0.0000 | 0.0000 | 0.0000 | 0.2430 |

**Table 2:** RegenPGC Segmentation Model: Classwise F1 scores for the evaluation set. F1 scores span the interval [0, 1] with values near 1 indicating near perfect precision and recall. * next to a loss function and bold rows indicates this loss was selected for baseline model training.

| Loss Function | Background | Quadrat | PGC Grass | PGC Clover | Broadleaf Weeds | Maize | Soybean | Other Veg. | Overall |
| --- | --- | --- | --- | --- | --- | --- | --- | --- | --- |
| **CE\*** | **0.9592** | **0.9326** | **0.6451** | **0.8374** | **0.7305** | **0.8037** | **0.0000** | **0.7972** | **0.8799** |
| *InvCE* | 0.9256 | 0.7566 | 0.4894 | 0.6279 | 0.3788 | 0.2814 | 0.0731 | 0.2962 | 0.7526 |
| *FL* | 0.9524 | 0.8775 | 0.6088 | 0.8253 | 0.6868 | 0.7050 | 0.0000 | 0.6730 | 0.8649 |
| *CB CE* | 0.9307 | 0.8308 | 0.4853 | 0.7339 | 0.4145 | 0.4462 | 0.1804 | 0.2472 | 0.7880 |
| *CB FL* | 0.7896 | 0.2806 | 0.4970 | 0.6094 | 0.3670 | 0.0934 | 0.0473 | 0.1368 | 0.6316 |
| **ACB CE\*** | **0.9578** | **0.9439** | **0.6457** | **0.8354** | **0.7255** | **0.7454** | **0.5441** | **0.7750** | **0.8776** |
| *ACB FL* | 0.9408 | 0.8101 | 0.5525 | 0.8055 | 0.6375 | 0.4673 | 0.2250 | 0.5740 | 0.8407 |
| **Recall CE\*** | **0.9491** | **0.9418** | **0.6518** | **0.8323** | **0.7334** | **0.8082** | **0.6493** | **0.7807** | **0.8686** |
| **Recall FL\*** | **0.9456** | **0.8802** | **0.6227** | **0.8184** | **0.6815** | **0.7325** | **0.5250** | **0.7343** | **0.8557** |
| *DL* | 0.9356 | 0.0000 | 0.5975 | 0.7769 | 0.0000 | 0.0000 | 0.0000 | 0.0000 | 0.8287 |
| *TL* | 0.9358 | 0.0000 | 0.5983 | 0.7786 | 0.0000 | 0.0000 | 0.0000 | 0.0000 | 0.8305 |

In comparison, losses which are designed to perform better than CE loss under unbalanced data conditions, tended to perform worse than CE in almost every class of of texture we modeled. InvCE loss had a 0.26 drop in the Jaccard index and a 0.12 drop in the F1 score when compared to vanilla CE. Similar results were seen for the models optimized using Focal loss and the CB class of losses. In particular, CB Focal loss had very poor performance, especially for the minority classes ‘maize’, ‘soybean’, and ‘other vegetation’ which is surprising given that this loss reweights and focuses the loss on specifically to target these minority classes. The Dice Loss and Tversky Loss scores, are both structurally different than many of the other loss functions which can integrate broadly with CE and Focal losses, and directly work towards optimizing a Jaccard Index or Intersection Over Union like score. However, both of these models only predicted pixels in the top 3 majority classes ‘background’, ‘pgc grass’, and ‘pgc clover’, and had a high degree of False negatives in every other class. The best loss classes were both in the Adaptive Class Balanced (ACB) and Recall class of loss weights, generally when used in conjunction with CE loss. ACB and Recall loss models converged to have an overall Jaccard Index of ≈ 0.60 to 0.65. These models predicted the ‘background’ and ‘quadrat’ classes with a high degree of accuracy and performed reasonably well on the ‘pgc clover’, ‘broadleaf weed’, and ‘maize’, but less than desirable accuracy in terms of ‘pgc grass’ when compared to the lower performing loss functions that had similar scores. From these results, we selected the top 4 losses to be used in baseline model training: CE Loss, ACB CE Loss, Recall CE Loss, Recall Focal Loss.

These results present an interesting set of data that need to be reasoned through. First, there are some classes, such as the background, which includes soil and crop residue, that are fairly homogeneous classes of image textures and really only vary in soil color across the training dataset. The same could be said for ‘maize’, where maize leaves are relatively distinct in appearance in comparison to other grasses in the image, even though they are in the minority of classes on the pixel class representation of the dataset. The ‘soybean’ class is has the lowest representation in dataset which makes sense why vanilla CE has a Jaccard Index of 0.0 for this class. While losses such as InvCE and CB losses were able to predict these pixels better, none surpassed 0.10 Jaccard Index or a 0.18 F1 score. In comparison, Recall loss did an excellent job at predicting soybeans (Jaccard Index=0.4807) by integrating the False Negative Rate into the network backpropagation.

In contrast to these relatively easy to predict classes, a wide variety of ‘pgc_grass’ appearances are represented, not only in terms of phenological manifestation (the growth patterns and morphological changes of grass throughout the growing season as shown in fig 4), but also in the number and variety of different cultivated and non-cultivated grass species showing up in the images. This presents a great classification challenge - even when this class is within the top 3 majority classes in the dataset, it does not have the third highest Jaccard index or F1 score for any of the models. In fact, it is usually has one of the lowest classwise evaluation scores when compared across all models. This highlights the unfortunate fact that more data doesn’t necessarily mean automatic increases in a classes prediction scores.

**Figure 4:**
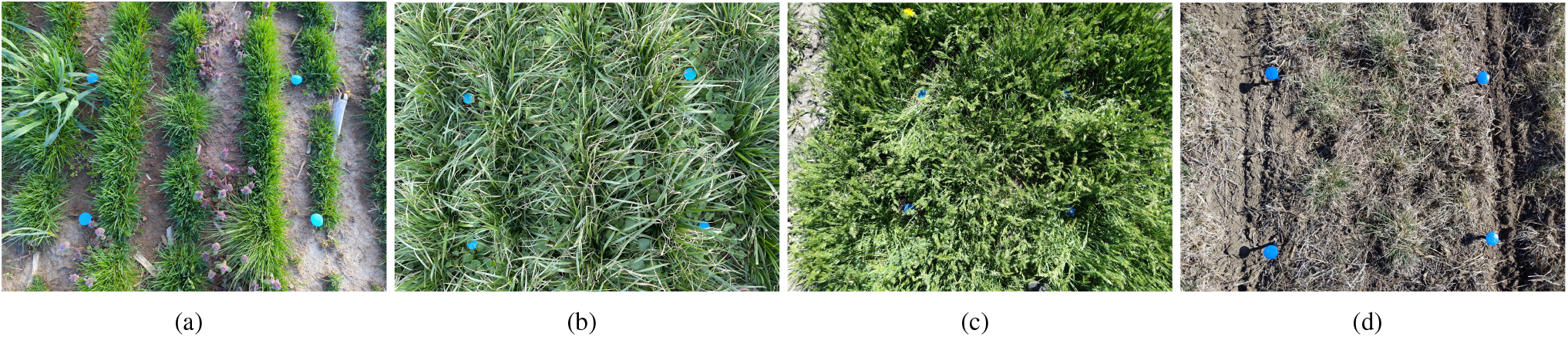
Examples of ‘pgc_grass’ in various phenological stages (a) PGC grass growing early in the season (b) mature stand mid-season (c) flowering stage (d) dormant stage.

Secondly, building off of the problem of classes with high appearance homogeneity and low variance across class instances, some of those classes, even when in the minority class, remain particularly hard to classify. The focal loss classes are designed to address this problem by focusing the weight on difficult to classify pixels, but this is highly dependent on the *γ* focusing parameter (eq. 4). Additionally, the CB losses are dependent on the hyperparameter *β* to calculate the effective sample number, and this has to be tuned to get decent performance. We performed no hyperparameter tuning due to time constraints, so we used the default values presented in the original manuscripts for each of these losses. Two of the benefits to using losses such as ACB and Recall Loss is that the reweighting procedure is not 1) arbitrarily calculated at the beginning of the the modeling process such as InvCE Loss, nor does it 2) require extensive hyperparameter tuning to get decent predictions from them. These loss weights are determined directly from the data and thus benefit from informed decisions during network backpropagation instead of making unreasonable assumptions about what the weights should be beforehand.

Thirdly, scaling down images with original resolutions of around 3000×4000 pixels to 512×512 drastically reduces the visual and gradient contrast between vegetation categories that are so distinct at higher resolutions. While this reduction was necessary to rapidly train several different models so loss criterion selection was possible, this presents additional challenges to the model, since many important features are blurred at such a low resolution. This is particularly evident when even trained experts have a hard time labeling messy images accurately at full resolution, let alone when the image size is reduced by 95%.

### 3.3. Baseline Model Training

Baseline weights using the top four loss functions were trained using the same DeepLabV3+ model decoder but instead with a more modern ‘efficientnet-b4’ backbone and a training and inference image size of 1024×1024. We instantiated the models using Vanilla CE, ACB CE Loss, Recall CE Loss, and Recall Focal Loss with standard stochastic gradient descent and trained using the parameters explained above in the approach. We saved the best weights from each training run based on the lowest validation loss and calculated the multiclass Jaccard Indices and the F1 Score. Overall the scores substantially increased compared to the preliminary models. Jaccard Indices increased by 0.1034, 0.1576, 0.1583, 0.1805 and the F1 Scores increased 0.0598, 0.0646, 0.0766, 0.0709 for CE, ACB CE, Recall CE, and Recall Focal Loss, respectively (table 3). These were simply the result of a better backbone and increasing the image size to 1024×1024. The only class that only saw a very slight increase upon model evaluation was the ‘soybean’ class when the loss was Vanilla CE. The ‘background’ prediction as measured by Jaccard index was almost perfect at 0.96 for all loss criterion. One of the largest jumps in Jaccard index and F1 score was seen for ‘pgc grass’ which was ≈ 0.45-0.48 in the smaller, preliminary models, but increased to 0.65-0.73 with these newly trained models. The same could be said for the ‘broadleaf weed’ category. Since Recall CE Loss had the lowest loss overall, we chose those as starting weights for the rest of our experiments.

**Table 3:**
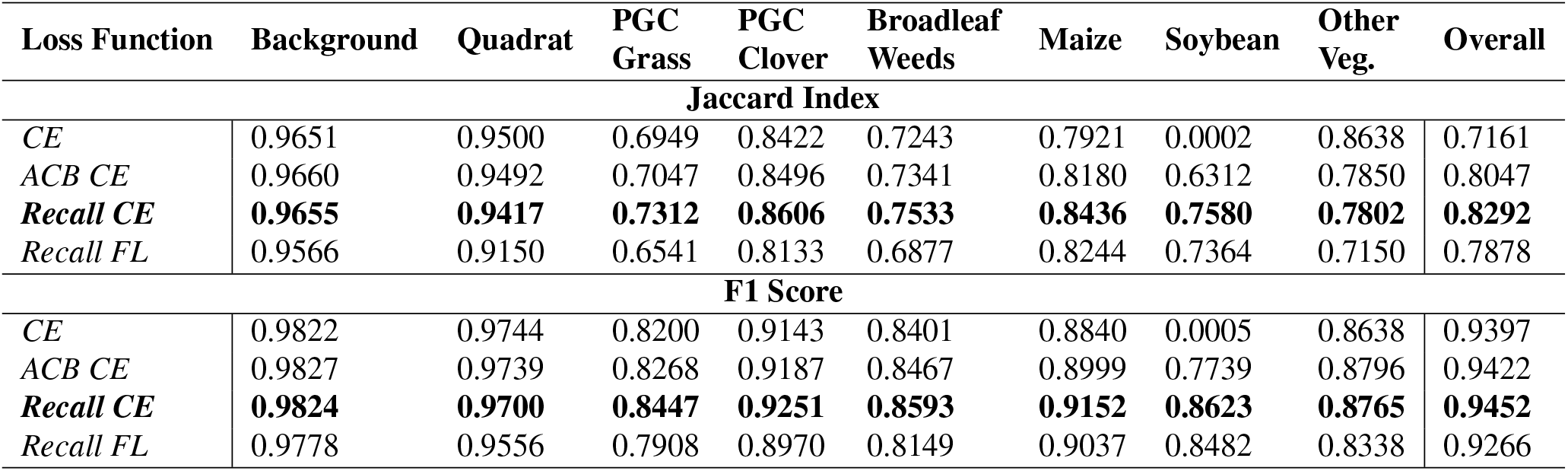
RegenPGC Segmentation Baseline Model Evaluation: Class-wise Jaccard Indices and F1 scores for the evaluation set. Both Jaccard Indices and F1 scores span the interval [0, 1] with values near 1 indicating more accurate predictions. Recall CE Loss was the best performing in most of the important categories.

The ‘other vegetation’ class was created to be able to model non-target plant distributions that appeared in one of the open-source databases but do not appear in any of the target RegenPGC images nor will they in the coming growing seasons. There were many broadleaf crops in this category that could easily have been mistaken for ‘broadleaf weeds’, but the overlap in predictions for these two classes was very low, ≈ *<* 1% of the images with ‘broadleaf weed’ labeled pixels had any of the pixels miscategorized. In fact, almost all of the ‘other vegetation’ predicted pixels occurred only in the images in the CAW dataset from which that class was wholly derived. One of the most fascinating aspects of these baseline models was the ability to take three vastly different image sets from different domains, combine them, and be able to train a single model that segments distinct classes found only one of the three datasets. Generally speaking, there was a high degree of correspondence between the ground-truth masks for any given training image, and the prediction mask the baseline model produced (Jaccard Index: 0.8292). This was true across all three datasets, as shown in fig 5. Still yet, these baseline models at this point were trained without any training input from the actual target RegenPGC dataset which includes ROI plot markers and vastly different grass species. In order to see how this current best model predicted on out-of-sample RegenPGC images, we ran through several examples and were surprised at the level of precision in the masks given that the RegenPGC images are in general much messier and more complex, than the oversimplified Kura Clover Dataset and the Crop and Weed Dataset (fig. 6, a-d).

**Figure 5:**
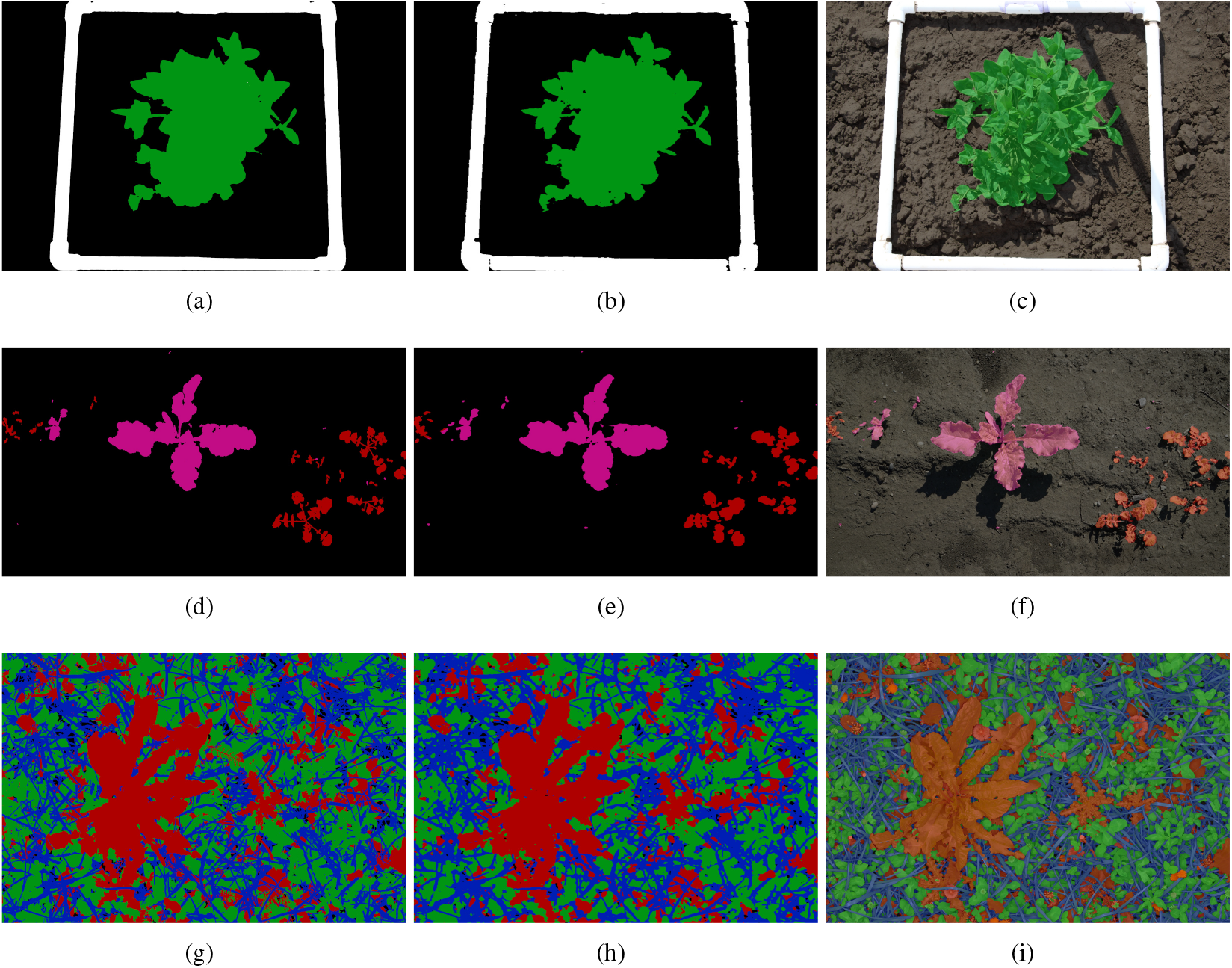
Inference examples using the baseline trained weights with Recall CE loss. All rows L to R represent the ground truth masks, model predictions, and the prediction overlay. Blue=PGC grass, Black=background, Green=PGC clover, Red=Broadleaf weeds, Pink=Other vegetation. (a-c) PGC clover image from our internal dataset. (d-f) CAW dataset inference example (g-i) GC dataset inference example

**Figure 6:**
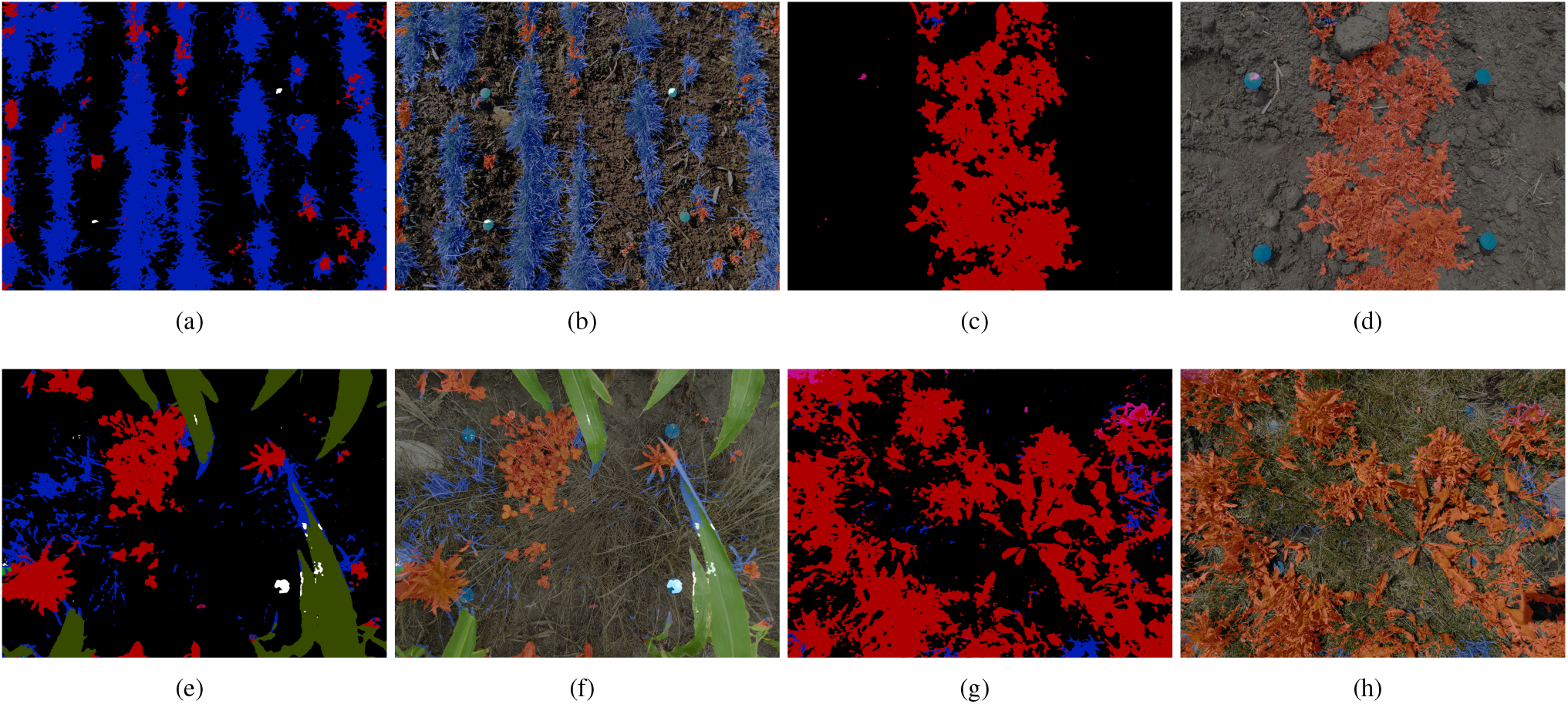
Inference examples using the baseline trained weights on out-of-sample, unlabeled RegenPGC images. While there is excellent correspondence between the predicted PGC grass (a) and the actual grass (b), and predicted weeds (c) and actual weeds (d), the model does not predict dormant grass well (e-h.)

One problem that was readily apparent was the inability of the model to classify dormant grass in the correct class. Every image that we ran inference over had dormant grass either misclassified as ‘background’ or as ‘broadleaf weeds’ which precludes the full use of the image analysis pipeline and prevents the green/brown pixel discrimination (fig 6 e-h). The main reason for this misclassification is simple - out of the 12k images used for training, there are no images with any brown/dormant grass instances. The closest class in appearance is, in general, the background class since the CAW, GC, and kura clover datasets all contain some level of crop residue (i.e. dried, dead plant material) left on the soil surface and these were all included in ‘background’. To combat this, we selected a total of 282 images directly from the RegenPGC dataset, which either 1) contained PGC grass in a different phenological state than the grasses in the CAW or GC dataset (flowering, dormant, etc.), or 2) contained a high mixture of broadleaf weeds and patches of soil interspersed with PGC grass, that was either dormant or active. We labeled the images quickly and then injected them into the training dataset to see how much improvement we would be able to get from a new model trained when we included the new data. We started training with the weights from the baseline Recall CE Loss model and kept the rest of the settings constant.

This improved model, what we will call the “injection” model, led to slight increases in Jaccard Index across all classes compared to the baseline RecallCE model. But most importantly increased the PGC grass Jaccard index improved to over 0.75 which we were not able to achieve with any of the other models. The injection model’s predictions for PGC grass show a lot of improvement over the previous iterations, especially in regard to the dormant grass which is an extremely important prediction category for the pipeline. Running inference on the same images from figure 6 above yields the correct predictions as shown in figure 7. The PGC grass mask predicted from this model covers all of the brown, dormant grass, and cleans up the masks and predictions from many of the other classes as well.

**Figure 7:**
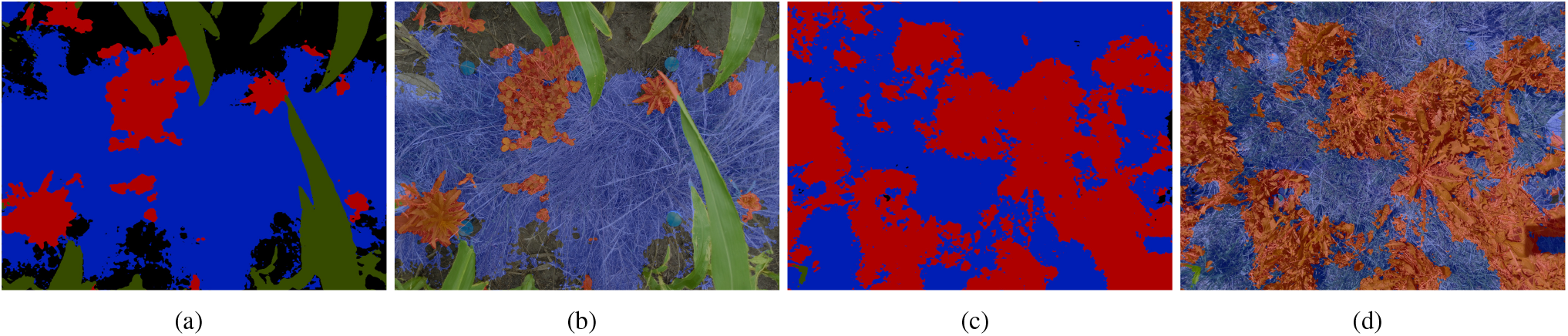
Inference examples using the injection baseline trained weights on out-of-sample, unlabeled RegenPGC images. Plot dormant PGC Grass prediction (b) Prediction overlay, (c-d) Image prediction and overlay

### 3.4. FixMatch Semi-supervised Learning

In order to implement the FixMatch SSL algorithm, we had to make significant changes to our Python trainer class that it subclassed. In brief, the FixtmatchTrainer class utilizes two separate Torch Datasets in tandem with different sampling strategies. The labeled dataset returns images and target masks as usual but the new unlabeled dataset loops over all of the images returning just the weak and strong transforms of the raw image tensor. Both of these datasets get sampled randomly. The longer dataset (generally, the unlabeled dataset) gets shuffled and sampled randomly throughout an epoch. However, since the labeled and unlabeled data are processed together in one batch step, both the labeled and unlabeled dataloaders must have the same iteration length. We solved this by implementing a customized infinite sampler for the labeled data such that while the unlabeled dataset is still iterating, the labeled dataset will continue looping over it’s shuffled elements until the epoch is complete. The two datasets return a batch of labeled data and a batch of unlabeled image transforms to be processed at each step. We started by concatenating the images into one tensor, passing through the model, and then returning the resulting logits to work with through the FixMatch algorithm we could save some computation (see figure 8a). This initial approach ended up being problematic for many reasons.

**Figure 8:**
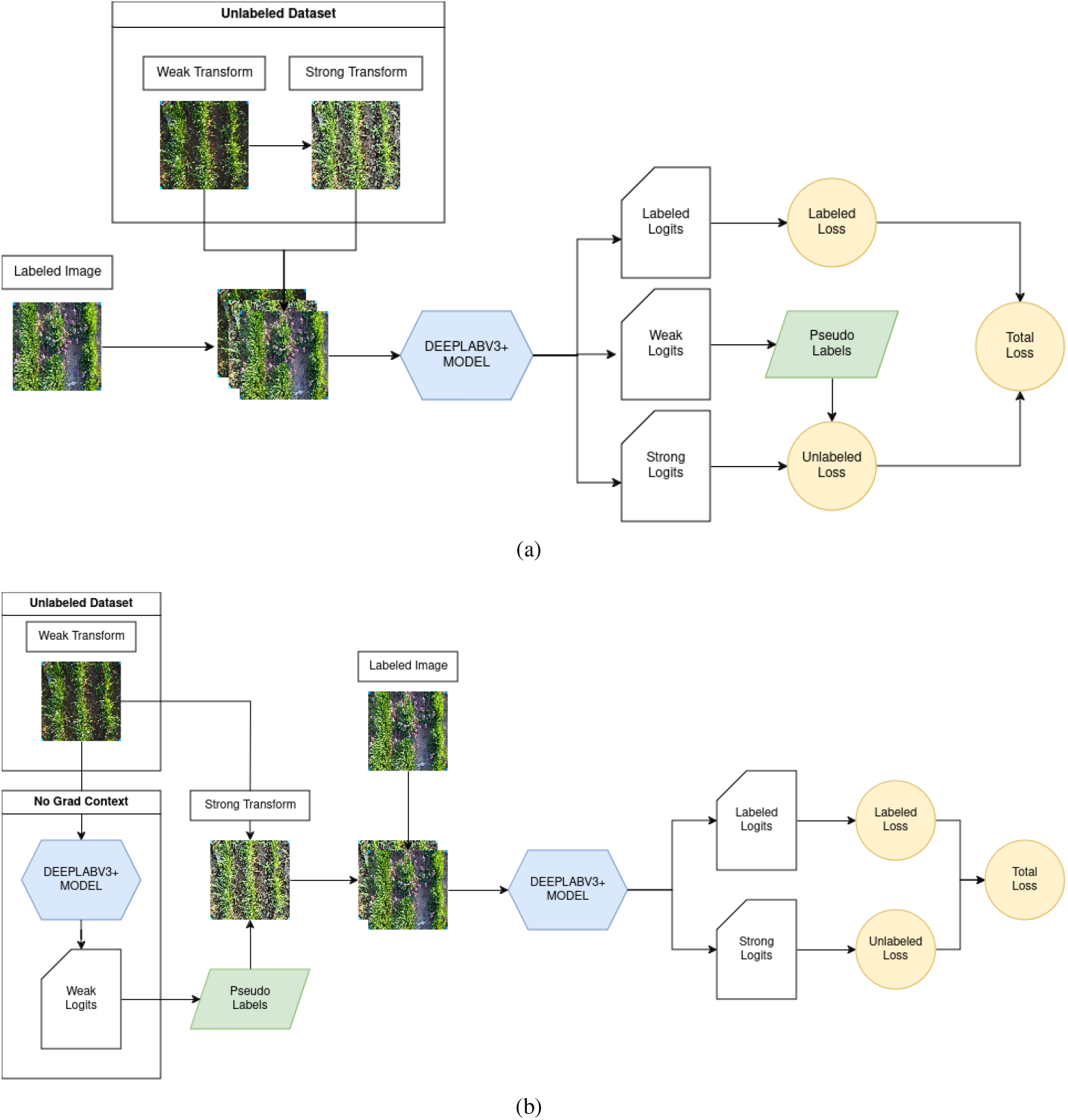
Our initial and final implementation of the FixMatch algorithm in Pytorch. (a) The initial but incorrect implementation of the algorithm. Strong transforms are applied within the dataset leaving no mapping between the strong transformed image and the weak psuedo-labels. (b) Correct implementation, weak logits are calculated without gradient accumulation and the strong transforms are conducted along side of the pseudo labels, constructing a 1:1 map to the output image. This require 2 model passes for each batch of labeled and unlabled images.

First, the image transformations for the unlabeled dataset were being calculated within the instantiated UnlabeledDataset class which precluded accurate calculations of the pseudo-labels for the strong images, since this class has no access to the model. Both the weak and strong augmentations include spatial transformations which rotate, shear, or perform affine shifts on the target masks. Initially, we passed the raw unlabeled image through the weak transforms, and then resulting “weak transformed” image to the strong transforms, finally returning both for each batch. Since each of 4 augmentations happened in the strong transforms with probability *p* = 0.5 and happen independently, we can expect to see spatially non-congruent regions between the weak and strong image with probability *p* = 0.9375, thus rendering the pseudolabels calculated on the weak images random noise since there was no longer any way to map between those targets and the strong transformed image.

Secondly, during the initial implementation, the labeled, weak, and strong images were concatenated and passed through the model once to get the resulting logits. These were split once returned: the labeled logits are used to calculate the labeled loss, weak logits are used to calculate pseudo-labels and confidence mask, and the strong logits are used in the unlabeled loss calculation. Due to the high chance of spatial transformation between the weak and strong transforms, the pseudo-labels were very likely incorrect. Furthermore, the weak transformed image takes no part in the loss calculations except for the fact that it generates the pseudo-labels. So passing this image through the model with gradient accumulation on, tracks all of the operations performed on that image, which could potentially lead to noisier updates. This image should have been passed through the model in a ‘torch.no_grad’ context such that those operations would not contribute to model updates during backpropagation.

We redesigned the implementation of the unlabeled dataset to only return the weak transformed image, pass that through the model once without tracking gradient operations, and then calculate the final pseudo-labels after passing the initial pseudo labels through the strong transforms with the image so the same operations get applied to them(figure 8b). Those labels and the weak image were then passed to a strong transformation function in the main training loop that performed the same transforms on both the target mask and the image thus mapping the pseudo-labels directly to the spatial layout of the strong transformed image. Then the strong transformed image and the labeled images were passed through the model at the same time, returning the logits used to calculate their respective portions of the total loss. Designining this took some extra time as there are many small details in the implementation of the algorithm that require attention to detail. Furthermore, we had initially designed this FixMatchTrainer class to be able to run either one one or multi-GPU settings using PyTorch Distributed Data Parallel (DDP) context. While running the fully supervised baseline models in DDP mode was simple to implement, we were not able get the FixMatch algorithm to run properly under this paradigm due to some errors with how the logit tensors were being collected from across GPU devices to calculate the different portions of the loss. Thus we were also limited in the total batch size we could run this training algorithm.

We tried several variations on this codebase once we fixed the design errors, but ultimately every single version yielded a set of model weights that performed worse than the fully supervised modeling schemes. Trying to tune this model was tricky as each training run took up to 3-4 days and so we could not quickly iterate through different hyperparameters to see what a feasible set of *τ*, *λ* and learning rates were. We also implemented the ability to handle a class-wise threshold *τ* so that we could selectively restrict the classifications on a class basis during pseudo-label construction. This would allow us to set a very high threshold for classes with high baseline prediction accuracy and lower thresholds for classes that were a bit difficult to classify instead of forcing all predictions through one threshold value. However, due to time constraints, we were not able to test this out. We trained the algorithm using *τ* values of 0.95, 0.85, and 0.75 but all of them yielded similarly poor prediction results. The closest we got to a better model was when we started out with the baseline RegenPGC “injection” model weights and trained for a total of 15 epochs and the confidence threshold *τ* = 0.75 to keep only relatively confident pixel classifications. This took roughly 4 days on an H100 GPU, and iterated over 12k images in the labeled training set and 36k images in the unlabeled dataset. The best set of weights from this were slightly better overall than the injection baseline when comparing the raw evaluation scores, but most of the predictions were actually slightly worse when compared to the baseline injection model. Specifically, the FixMatch weights had a tendency to grossly underestimate the extent of PGC grass in the image and instead classified it background or weeds. Our worst FixMatch model ended up with exploding gradients when we started model training from randomly initialized weights. We tracked the proportion of confident pixels in each batch to see how this changed. This proportion oscillated wildly through each batch until eventually the gradients blew up causing the training to fail. Since the success of the FixMatch algorithm relies heavily on the concept of ‘consistency regularization’ between the weak and strong image transforms, it is possible that the strong transforms augmented the image too much, precluding the model from making correct predictions, i.e. over regularizing the model. We tweaked the strong transforms into a more reasonable range, and retrained, but this had little to no effect on the overall model performance.

Having a large, diverse set of images helped in the initial baseline model training, but for Fixmatch it’s possible that the out of sample images are hurting rather than helping refine the weights since we tended to have worse predictions once we added in the larger set of RegenPGC images. We tried one last approach that seemed reasonable. First, we defined a new labeled training set that consisted of the 282 labeled RegenPGC Images, and then added a total of 100 random images from the GC dataset, 30 images from CAW that contained corn, and 90 images from the internal Kura Clover dataset. These 502 images in total comprised the new labeled training set. We then defined the unlabeled training set as a total of 4794 images which included an additional 100 images from GC, 90 images from the Kura Clover dataset, an additional 30 images from the CAW containing corn. This new rebalancing was done 1) to reduce the total number of images in the training data so that we had a shorter cycling time in training iterations, and 2) so that the vast majority of images in the training set were in-sample target dataset images. We ran the FixMatch training scheme using *τ* = 0.75 and kept all other parameters the same as the previous FixMatch training attempts for 25 epochs. The model was initialized with the “injection” model baseline weights. After just a few epochs, the prediction outputs were remarkably better when performing inference on the unlabeled RegenPGC images. However, since the original validation dataset, which contained mostly images from CAW, and GC was no longer as relevant to the predictions, validation performance decreased slightly. While we do not have a good estimate on a true “in-sample” validation set, the PGC-grass Jaccard Index increased to 0.83 for the labeled training data, and the F1-score for this class also increased to 0.90. Some comparison inference between the best, fully supervised model weights and these new FixMatch weights show more accurate predictions particularly around the soil, weed and PGC-grass boundaries as seen in figure 9. While this improved set of weights appear to be a step in the right direction, there is more work to be done, particularly in the weed and clover class discrimination as seen in 9b. This, however is understandable since, many wild clover species appear in plots alongside of weeds.

**Figure 9:**
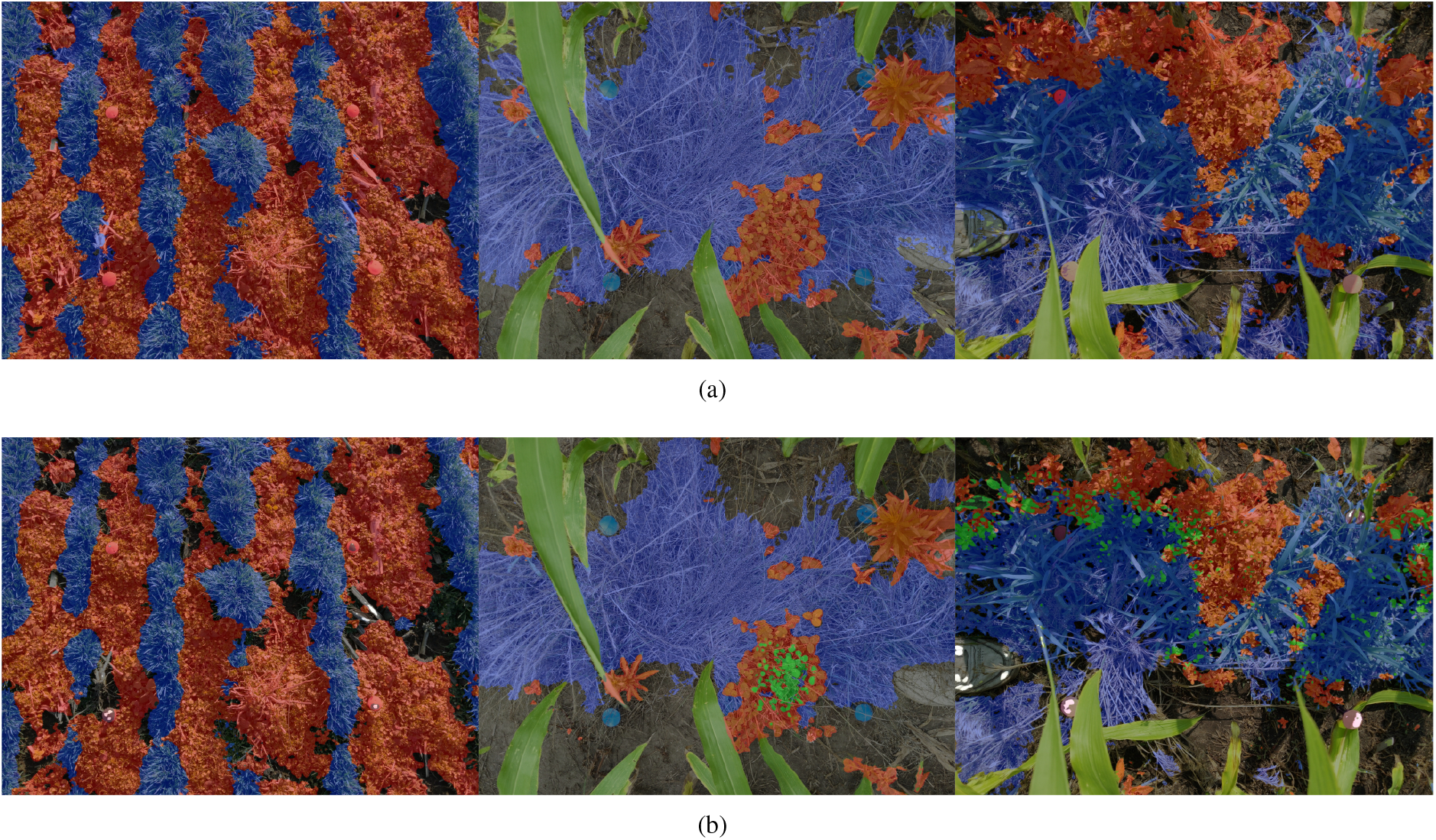
Baseline injection predictions compared to improved FixMatch predictions when trained on a focused subset of data (a) L to R model mask predictions on 3 representative images using the baseline model “injection” weights. Weed and grass masks tend to overestimate the extent of their boundaries, encroaching onto bare soil. Wild clover in the middle picture is incorrectly classified as weeds. (b) Improved FixMatch model weight predictions. Weed and grass masks are much tighter and the soil/background class is better accounted for. Wild clover in middle image is partly classified correctly representing a step in the right direction. Some issues with weed/clover discrimination on the right-most image indicates more training examples are needed to better discriminate between these two classes.

Semi-supervised learning (SSL) is an open research area that has tremendous application potential when applied correctly. However, it works best when performed on balanced datasets, so applying it to our taks was difficult, especially when rebalancing methods rely on accurate counts of the data classes. In SSL, these true values are, by definition, unknown. We suspect that our initial implementation did not do well for two reasons. First, since the labeled training datasets are different in composition and label structure, there is reason to believe that there is enough difference in the labeled and unlabeled data distributions to preclude efficient implementation of the FixMatch algorithm. In our first implementations, we included very few of the target images in the labeled training data, so the representations of each class which the model learned were slightly different than the unlabeled RegenPGC data. While this did not affect the fully supervised model training, it tended to make the predictions in the FixMatch training scheme worse due to non-confident predictions. Secondly, it seems that starting with a reasonable set of initial model weights helps model convergence. When we started training from randomly initialized weights, there were significant problems with convergence, and the model losses oscillated wildly out of control. In our final FixMatch implementation, we subset the full dataset to include a much higher proportion of the target dataset in both the labeled and unlabeled data while decreasing the overall size of both datasets. While this qualitatively improved the accuracy of the predictions, it is difficult to quantitatively prove this without a better validation set. In the absence of a large portion of labeled target data, removing labeled images from the training set to make a validation set can be a costly decision. Clearly, the absence of labeled data is not as big of a problem under and SSL training scheme, but validation of SSL models can be more difficult. This is something that needs to be addressed in future versions of the segmentation model.

### 3.5. Model Interpretability

While evaluating the predictions for both the baseline “injection” model and the FixMatch models, we noticed another startling problem. Whenever there was a sampling quadrat somewhere in the image, the FixMatch model had a high propensity to classify any weeds or grass in the image as ‘pgc_clover’ even in scenarios where very similar images without a sampling quadrat had been correctly classified. We reran inference on all the images using the baseline-injection model and noticed the exact same thing (figure 10). Unfortunately this is due to an oversight one our end when building the dataset. Out of the labeled training data (12643 images), 816 images contain a quadrat. Out of those, 800 have clover and the quadrat co-occuring.

**Figure 10:**
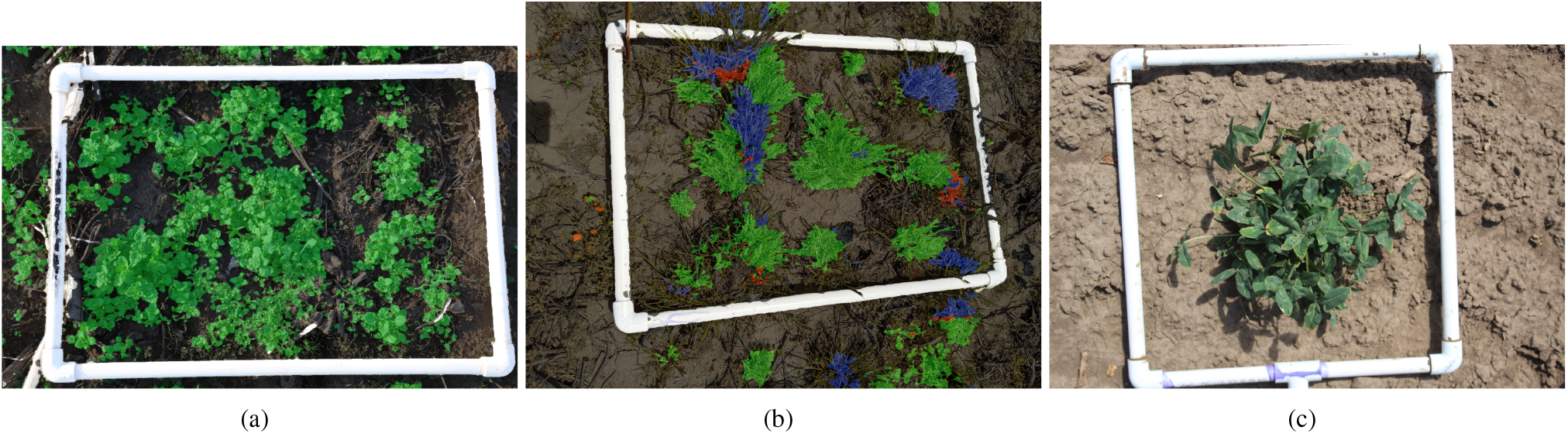
Incorrect predictions involving a quadrat when (a) broadleaf weeds are misclassified as clover (green overlay) and (b) PGC grass is misclassified as clover. (c) A representative image of a training image with kura clover inside a PVC quadrat and no vegetation outside. Notice also how there is vegetation outside of the quadrat in inset (a) that has been misclassified as ‘background’

The presence of clover is in 6,577 images, so a little over half of the images. If we define *A* = “Clover” and *B* = “Quadrat” we can easily calculate the conditional probabilities *P* (*B* | *A*) = 0.1216 whereas *P* (*A* | *B*) = 0.9814. These probabilities were mirrored when we ran inference on just the RegenPGC unlabeled images. The marginal probability of clover in any image was low, *P* (*A*) = 0.0343. The same could be said for the chances of seeing a quadrat, *P* (*B*) = 0.0444, and the marginal joint probability is *P* (*A* ∩ *B*) = 0.0292 which leads to an observed conditional probability *P* (*A* | *B*) = 0.6577. When we compare this to marginal probability of *A*, the *z* statistic is roughly 180, and p-value is essentially 0. So the prediction of vegetation as ‘clover’ upon inference is highly correlated with the actual appearance of a sampling quadrat in the image and not necessarily upon the actual presence of clover in the image. This is further corroborated by the fact that there is no intentionally planted clover in any of these unlabeled images and thus the *P* (*A*) is probably even lower than what is presented. We quickly implemented a class activation map visualization using Grad-CAM [52] of some of the salient classes for the final convoulutional layer of the best FixMatch model to see which areas of the image are most influential overall for each class’s predictions (figure 11). While this was a rough implementation, it shows that for some of the classes, the final classifications may not always be based on what you would think are the most important features for classifying pixels in a particularly category. For instance, in figure 11a, we can see that when classifying a quadrat, the quadrat areas are indeed highlighted over other areas of the image, but the important features related to the clover classification have almost nothing to do with the clover itself, but instead rely on the presence of soil or other objects in the image 11b.

**Figure 11:**
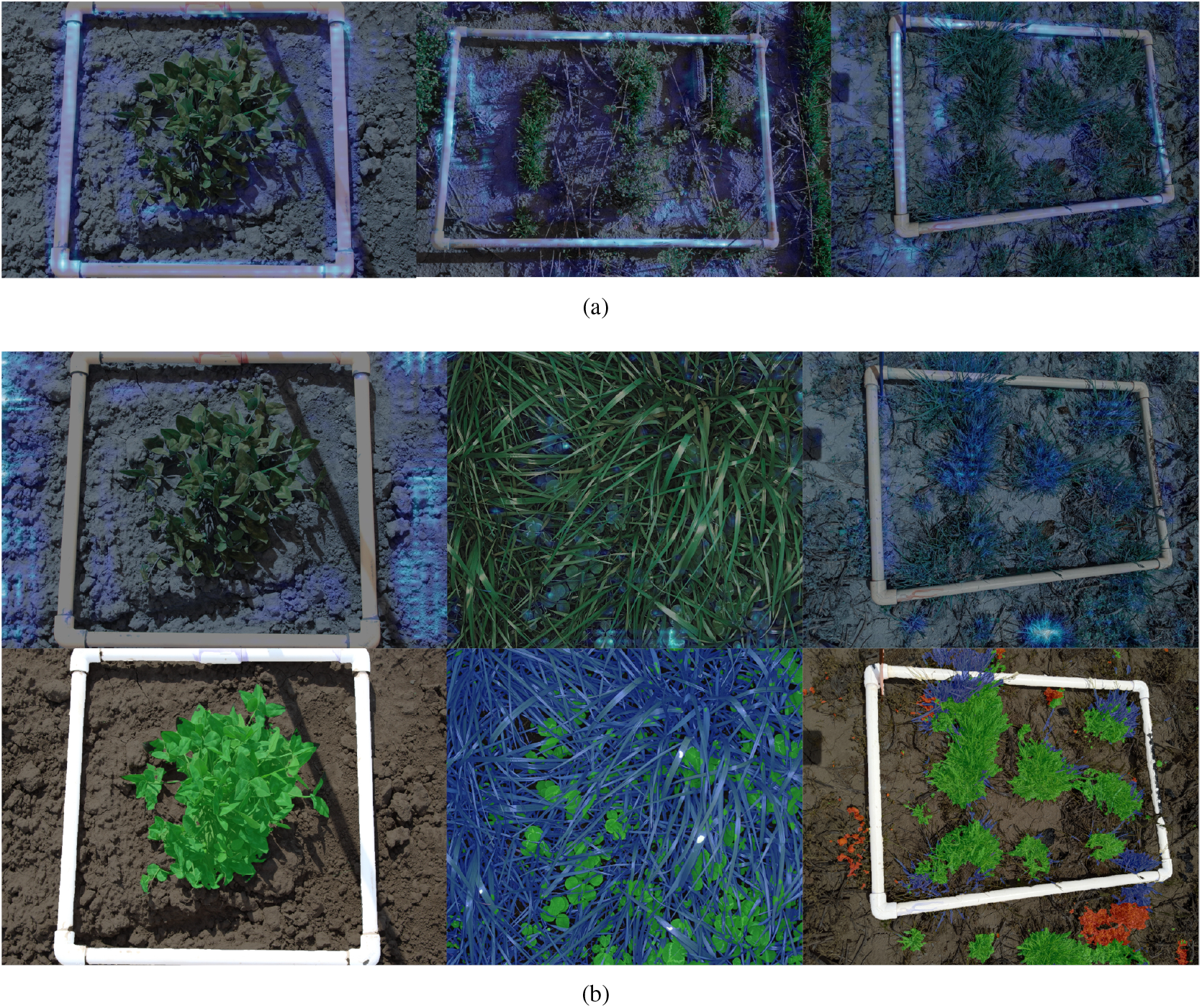
Grad-CAM Class activation maps to highlight salient features of the image that contribute overall to a class’s decision boundary. Light blue areas indicate strong activation for a class, darker areas indicate portions that do not contribute to a category’s classification (a) Grad-CAM applied to the ‘quadrat’ activations. This correctly highlights portions of the quadrat. (b) Grad-CAM applied to the ‘pgc-clover’ class. Top Row: Grad-CAM maps. Bottom Row: Prediction overlays. From left to right. Left) Grad-CAM highlights the soil around the quadrat and clover, but does not highlight the clover itself as contributing to the classifications. Mid) Clover in the Grad-CAM map is highlighted over the clover and the model correctly predicts the clover. Right) Grad-CAM shows that the clumps of grass are highly activated areas for the clover classification even though no clover is present. The prediction map show most of the PGC grass mistakenly classified as clover

While the Grad-CAM maps show some discrepancy in how the clover class is classified, It is still difficult to interpret them due to the fact that in some cases the model correctly identifies clover based on the activations of the clover, in other cases, based on the soil. In the third case, based upon the presence of PGC grass. While we don’t at this time fully understand the nature of these decisions, this warrants further investigation. In particular, we think that since there are few images of clover alongside of grass and weeds, this class would benefit from further refinement by including better training and validation examples or a more diverse set of training images. It is also relevant to consider that classes in the dataset should be more independent of one another, clearly not the case for the ‘pgc_clover’ and ‘quadrat’ classes. While not all researchers will be using sampling quadrats instead of markers, it is a fundamental flaw of the model if it classifies all vegetation in the model as the wrong class because of learned correlations in the training data rather than learned representations of a given vegetation class. As this project is ongoing, this is clearly an area of the dataset architecture that will be improved upon in future model versions.

### 3.6. Image Analysis Pipeline

The RegenPGC View image analysis pipeline draws together the marker detection model and the PGC segmentation model and some morphological image analysis techniques in order to extract the data that we really want, namely the proportion of pixels which belong to each class in the ROI, and the proportion of PGC grass predicted pixels that are green and active versus the proportion that is brown and dormant. This pipeline was explained in detail in 2.7 so we will just go over the main results. The first step in the pipeline is getting the image in the proper format since both models require the image to be 1024×1024 and normalized appropriately. Locally, the image just gets read in, resized and normalized, converted to a tensor and sent directly to the model for the forward pass. The same image gets then passed through the PGC segmentation model to get the raw logits. In production, the raw image gets read in and resized to 1024×1024 locally. Then we serialize the image in ‘application/x-npy’ format, and send to the AWS Sagemaker endpoint for model inference. Once in the Sagemaker instance, the image is reshaped and normalized before being sent to the model for inference. Part of the reason for splitting the operational responsibilities between local and cloud resources has to do with the size of normalized 32bit float images vs 8bit unsigned integer images when sending over the API as Sagemaker serverless endpoints have a strict 4MB payload limit. Whether in the cloud or locally, both models return the same data. The segmentation data gets processed to construct the class map as described above and the bounding box coordinates get returned and filtered for the top 4 detections.

Once we have the midpoints of marker, whether they be the quadrat corner or a landscape fabric stake, we need to find the transform matrix *M* which describes the mapping from a set of four source points to a set of 4 destination points, possibly on a different scale. While this is easy to solve in many software systems by solving a system of [linear equations](https://github.com/opencv/opencv/blob/11b020b9f9e111bddd40bffe3b1759aa02d966f0/modules/imgproc/src/imgwarp.cpp), the challenge can determining which marker in the ROI corresponds to which corner of the resultant image. Trivial solutions to this exist when the ROI is relatively square compared to the original image (see [here](https://pyimagesearch.com/2014/08/25/4-point-opencv-getperspective-transform-example/) for an example), but this does not always work and can sometimes lead to incorrect ordering or index errors. However, if we construct an L2 norm distance matrix between each of the points in the marker predictions with the image corners, we can use the linear sum assignment algorithm which minimizes the total cost of an ordering between the rows and the columns of a cost matrix *C*, such that we *minimize* 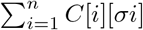 where *σ* represents some permutation of *i, C*_*i,j*_ represents the assignment of row *i* is assigned to column *j*, and each row is matched to exactly one column, in a way that reduces the overall cost of the weighted sum. If we think about the euclidean distance as the cost, this makes the process of matching the ROI corners to their proper image corner trivial and exact. We can then use this ordering to calculate the transformation matrix and return just the ROI from the image as shown in figure 12.

**Figure 12:**
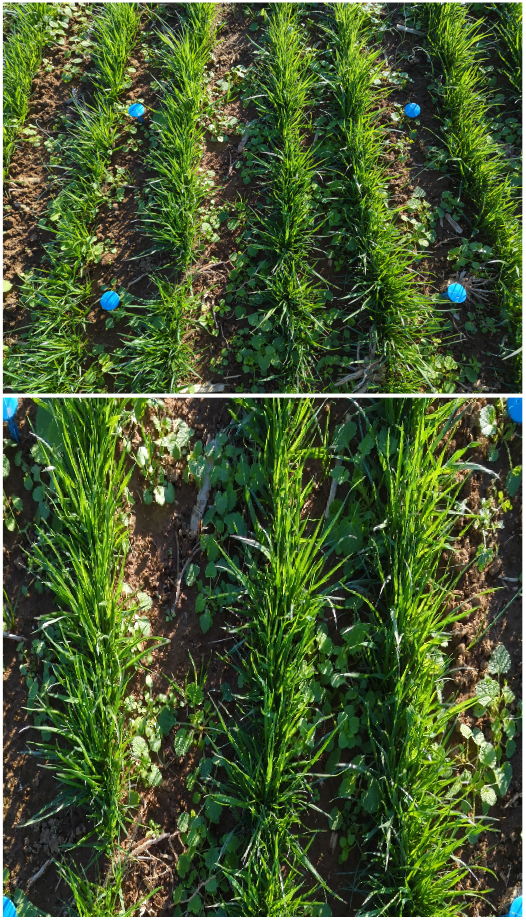
ROI transform from markers to standardized image

To calculate the proportion of pixels in the ROI that belong in each class we simply apply the transformation *M* to the mask, sum the non-zero pixels across each channel and divide by the total pixels in the image. These values are stored in a pandas DataFrame with appropriately named columns. For the green/brown pixel discrimination, we need to find a threshold between their values that we can use to separate them. We tried both a Gaussian Mixture Model approach with and without SLIC as well as a naive thresholding approach in the HSV color space. The ROI image was first converted the HSV and then we vectorized the image to be in ℝ(*m×n*)*×*3. Using a set of green/brown range mean values, we set the number of Gaussians to 2 and then used these means as prior means before iteration. We also tried a different version of this by preprocessing the image using SLIC to generate superpixels from the image. Both of these attempts failed due to a few reasons. First when using SLIC, one of the main assumptions for the algorithm is that there are large patches of homogeneous color and texture that can be simplified by creating one superpixel representation of it. This is clearly not the case as there is a high amount of variation just between the blades of grass in the ROI. So SLIC took areas of high entropy and information, and made them all one color. In fact, in areas where there was a lot of brown and green grass together, it coalesced into one superpixel with an RGB value somewhere between green and brown, and was not useful for discrimination purposes. Additionally, using SLIC was extremely slow and was, practically speaking, infeasible to use during inference. Second, the Gaussian mixtures even when setting prior means for each cluster, tended to drift away to find the most natural Gaussian clusters in the data, not necessarily the clusters that corresponded to the green and brown range on the HSV color scale. There was also an issue that when portions of blue markers and quadrats are in the ROI transformed image. These outlier colors tended to throw off the Gaussian calculations, as they were drastically different from what either green or brown Gaussian model would naturally predict, and so threw off the means and variances of each generative model.

For these reasons, the two approaches above were abandoned for a much simpler approach based on HSV thresholding. We were able to find a set of high and low HSV threshold values that we could pass the grass mask through and obtain the green and brown rage of pixels easily. When the contrast between areas of deep shade, and intense sunlight made the contrast in the image difficult to work with, we converted the image to the l*a*b* color space, split off the l* channel, and applied the CLAHE algorithm to even out the lighting across shade and sun. While this lightened the images overall, it had very little effect on the thresholds other than to make it easier to discriminate between green and brown in areas of intense light or shade (see figure 13). Once we have the transformed ROI class masks it is a simple masking procedure to extract just the predicted PGC Grass from the ROI and then apply the HSV pass threshold parameters to generate the proportions of active grass pixels and dormant grass pixels within the ROI. A simple extension of this is to use each mask to generate RGB images which include just the brown or green grass within an ROI (figure 14a). Finally, using the same four point transformation approach used to crop to the ROI, we can back transform the ROI mask to the original image to overlay whatever pattern (RGB prediction overlay, PGC grass masks, etc) to the original image (figure 16). While none of these visualizations are necessary to extract the needed data, they are good ways to debug any problems that are not readily evident from the data. For instance, out of 2500 test inference images there were about 10 with poorly transformed ROIs due to erroneous detections with the marker detection model which can be seen in figure 14b.

**Figure 13:**
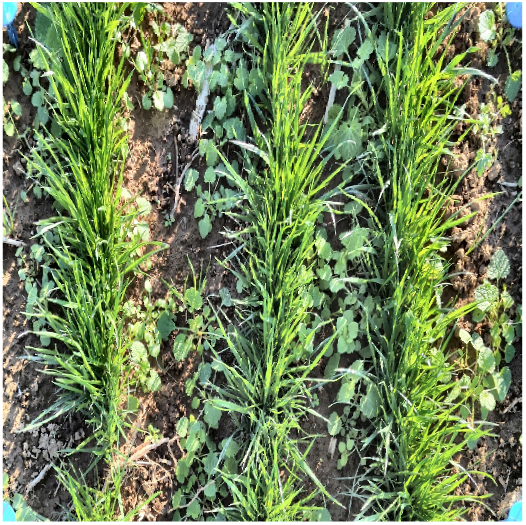
CLAHE algorithm applied to lightness channel of ROI transformed.

**Figure 14:**
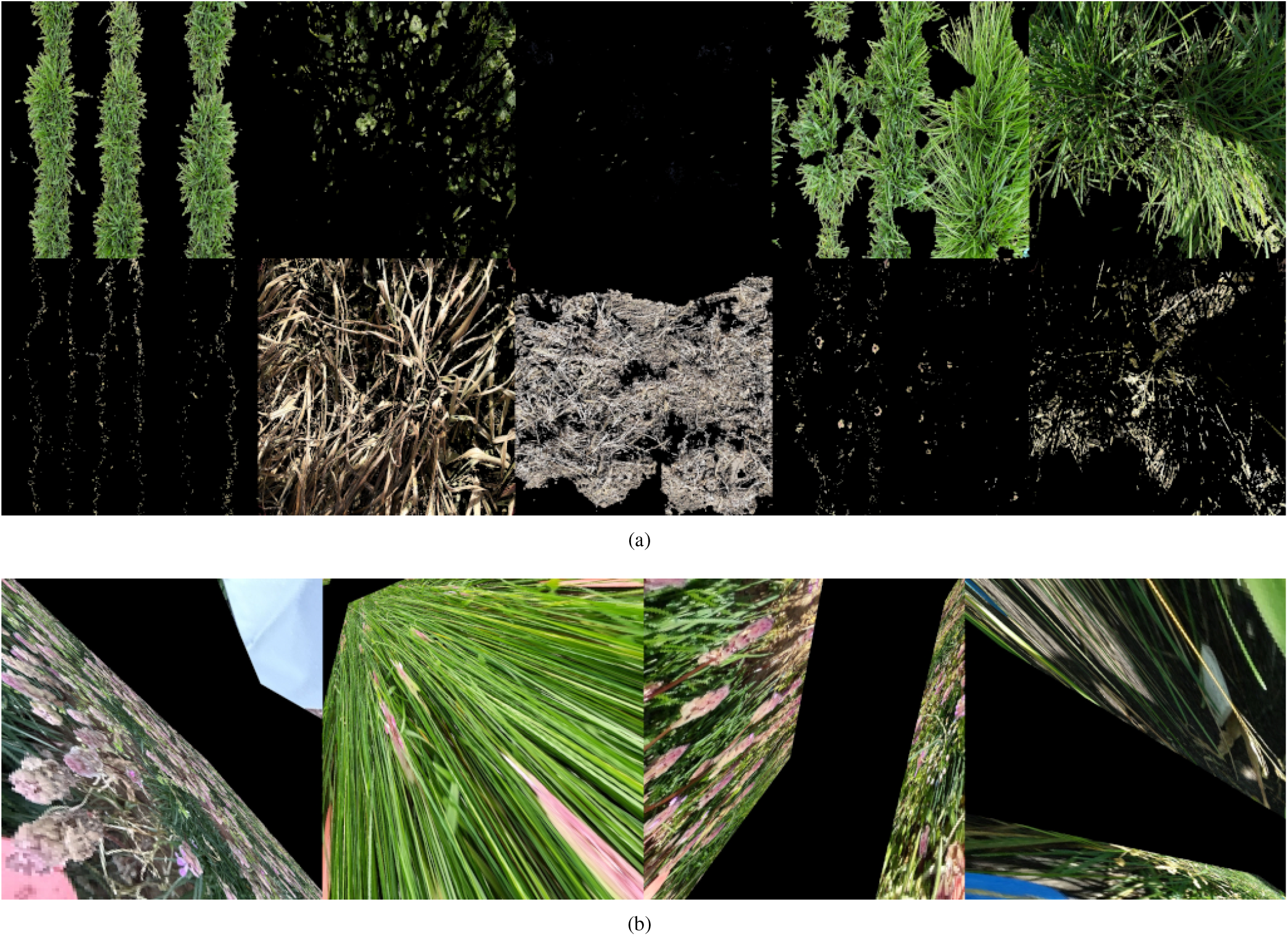
ROI transformations possible (a) Active and dormant PGC grass discrimination applied to transformed ROI grass masks. Top row: active portion of the predicted PGC grass pixels. Bottom row: Corresponding dormant portion of PGC grass pixels for each ROI. (b) Anomalous ROI transformations dues to erroneous marker detections. These will have to be corrected manually

**Figure 15:**
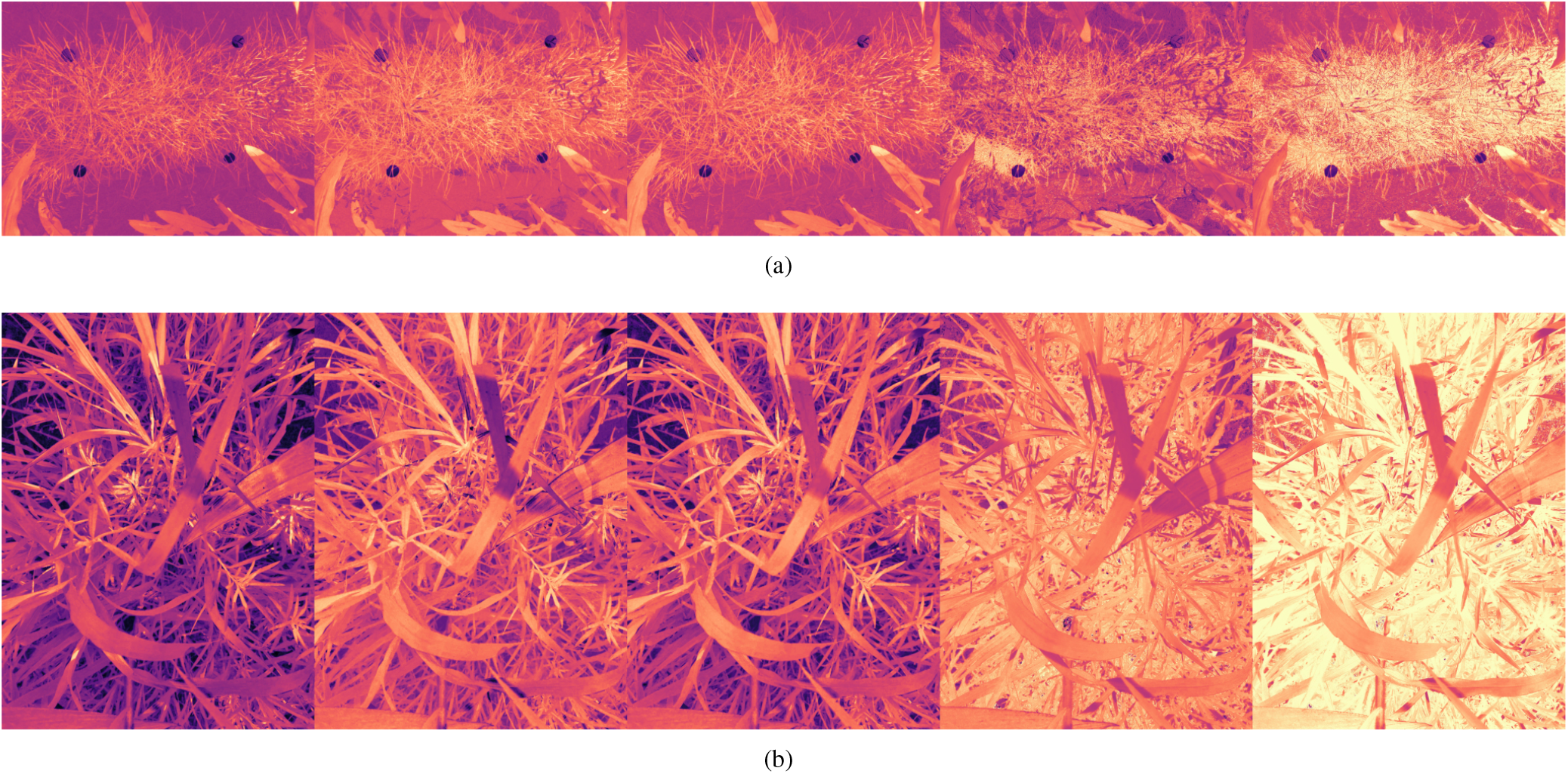
RGB vegetation indices pseudo-colored to indicate value. Warmer colors indicate higher values i.e. greener, healthier vegetation. From L to R: CIVE, ExG - ExR, MEXG, MGRVI, RGBVI (a) VIs applied to example a. (b) Applied to example b

**Figure 16:**
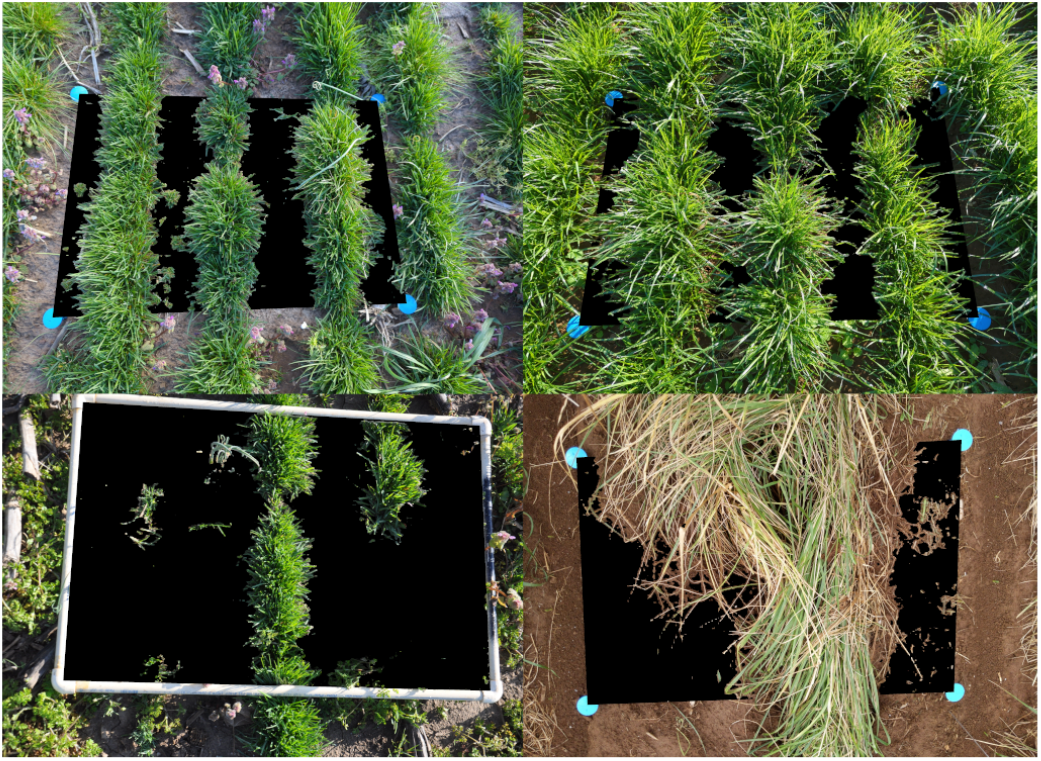
PGC grass prediction mask within the ROI back transformed to the original image.

As a part of the pipeline we evaluated 18 different RGB vegetation indices that can be used to distill RGB values down to easier to interpret scalar values for each pixel. Many of the indices just use linear combinations of RGB values such as Excess Green minus Excess Red (ExG-ExR) [36] or Color Index of Vegetation (CIVE) [27]. These tend to be the most resistant to noise and require very little processing of the data as they have set upper and lower bounds that can easily be transformed to a [0, 1] interval. Others such as VARI [22] are normalized values and can produce excellent, intuitive results depending on the right circumstances. However, they require value clipping to avoid ‘Nan’ values or extreme deviations from the normal range of the index and this requires extensive trial and error to determine the best range to restrict the values to without losing important information. Currently our pipeline has a few of the more linear VI’s implemented in their final form, but the others require more testing before they can be fully integrated. RGB VIs are inherently noisier than the more common NIR-RGB indices since the NIR captures a much better estimate of the health of plants reflecting light in the NIR range. RGB indices try to replicate these results by using a combination the RGB channels instead, so are inherently more limited. Furthermore, uneven lighting conditions can wildly affect the values of many of these VIs so care must be taken when acquiring photos. While the indices are interesting in and of themselves as an extra data point for the PGC grass, we performed some tests on these to see if the range of VI values could also be used to discriminate between active and dormant grass pixels within the ROI. Briefly, each VI was scaled down to [0, 1] and then we tried to find a single cutoff threshold below and above which we could construct dormant and active PGC grass masks. This was possible and when we used some of the VIs such as MEXG and Exg-Exr we were able to achieve decent results that rivaled the HSV thresholding approach. However, in many images, the lighting conditions were poor and the resulting active/dormant PGC grass masks were insufficient. One alternative to using just one or the other is to add the VI as a channel in the image and perform the thresholding over 4 channels instead of 3. We are actively looking into this avenue.

### 3.7. RegenPGC View Web Application

We had set out to develop a simple web application that would accept image uploads and run inference them and process them through the RegenPGC View image analysis pipeline. This application was a reach goal for this semester and is currently still under development and should be completed in the coming months. Unfortunately due to the shortened length of the summer semester, we did not have enough time to bring it to completion. Currently the main pipeline code is residing in the application but is not hooked up to the front end as of yet. We have some main application functionality built out though. First we have a functioning user registration page where people can register for a new account or reset their password (fig 17a) Currently this will be restricted to people with certain institutional domains so that only select people will be able to work with our system. Second, users can login and access their own homepage (fig 17b). The application will restrict their content to inference jobs that only they have submitted. Third, users can upload images and create a new inference job (fig 17c). The job creation page accepts multi file uploads and saves images to the server local storage in a timestamped folder with the job name. The application also extracts image metadata and persists both the job and the image metadata to the PostgreSQL RDBMS (fig 17d). Once users submit the job, and the data is persisted to the database, this will start a series of async jobs where images will be sent to our model endpoints in AWS Sagemaker one at a time, and then return the inference results. These results (bounding box coordinates, prediction masks, etc.) will be processed by the RegenPGC View pipeline locally within the app and the resulting plot ROI estimates will be persisted to the database with a primary key of the image id and the job id. Simple inference jobs with a few images will likely be done within a few seconds, but jobs with many images will take some time. Users can check the job status page for more information regarding one of their submitted jobs (fig 17e). When a job is completed and selected the user will be able to see a table version of the their results and be presented with an option to download the data as a csv file as well as some of the image intermediates. While this portion of the project was not able to be finished, we look forward to the potential of having such a system built out and fully operational.

**Figure 17:**
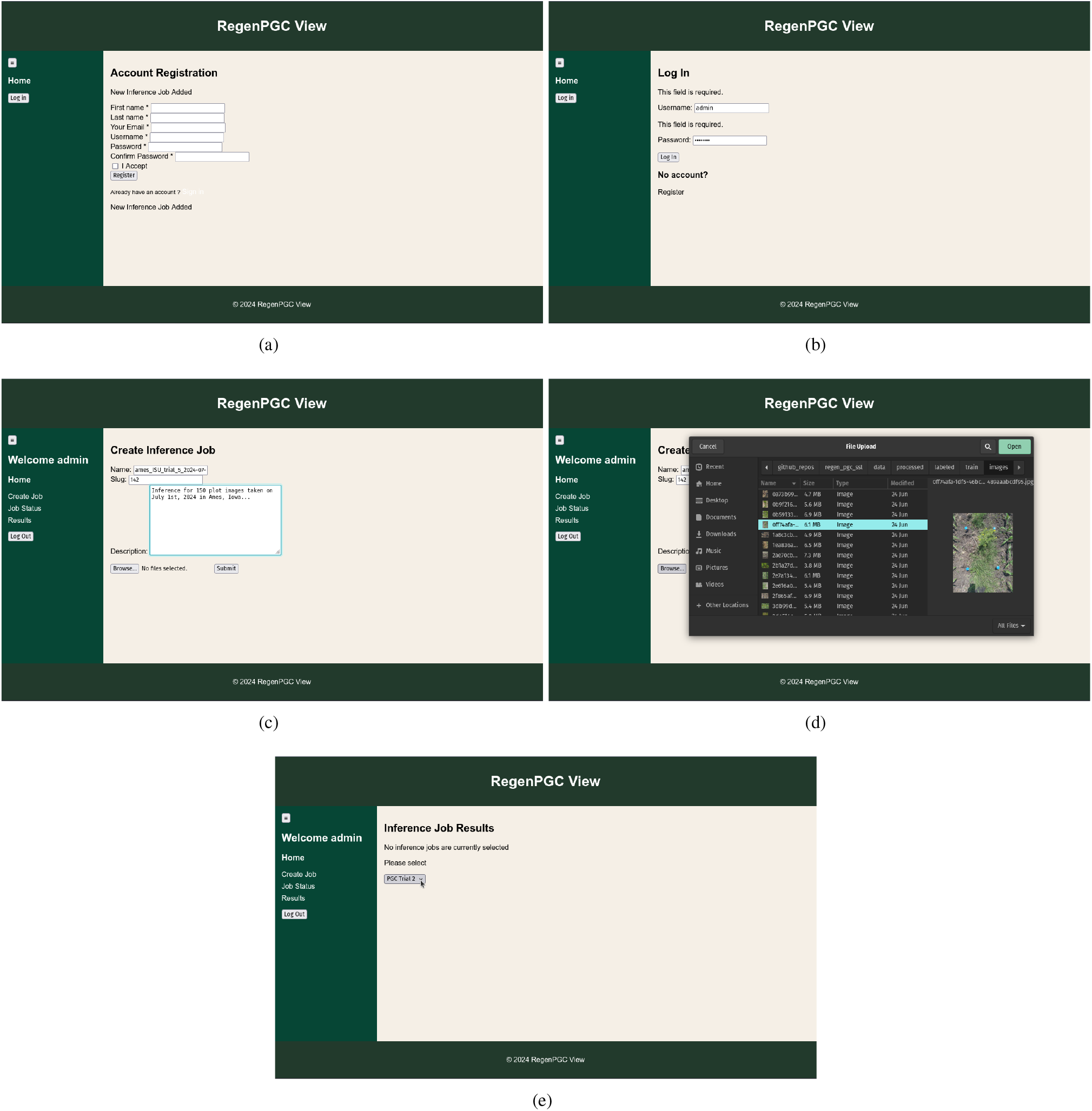
The RegenPGC View web application currently under development (a) Account registration page (b) Login page (c) Creating a new inference job and setting the job description (d) Multiple file upload to application (e) Checking the status of a current or past inference job

## 4. Conclusion

Perennial groundcovers are a novel, promising way to integrate cover cropping into conventional row cropping systems, all while minimizing soil erosion and runoff. However, due to methodological reasons it is difficult to quickly and objectively measure or estimate how well PGC plots have grown and established. In this project we developed a deep learning pipeline dubbed ‘RegenPGC View’ that leveraged both open-source and internally developed datasets to detect plot ROI corners using two different methods, segment each plot image into its respective vegetation/object class, transform the image to just the ROI to calculate the exact proportions of each class pixels, determine the number of green and brown pixels in the predicted grass mask, and then output all the data into an easy to consume format such as a csv file or push the results directly into a database. All of these steps were tied together into one pipeline to run sequentially. Our image detection model achieved near zero loss values and detection and classification accuracies above 95% for all classes. The PGC segmentation model achieved a high overall Jaccard Index and F1 score, and performed well on a class by class basis, but could benefit from further refinement. Our baseline, fully supervised models were able to achieve high prediction accuracy and overall Jaccard Index even though they were trained on related, but non-target, image data that has relatively little resemblance to our target RegenPGC dataset. When we included a few hundred labeled samples from our target dataset in the training data and trained this ‘injection’ baseline model, we saw some significant increases in the model performance though it was still prone to misclassification errors. While the FixMatch Semi-supervised learning portion of the project did not proceed as we had originally planned with the full dataset, we were able to train a much more performant model than the best fully supervised ‘injection’ baseline model when we restricted the labeled and unlabeled data to a majority of our target RegenPGC dataset images. However, the lack of more labeled data makes validating this model difficult, thus we relied on more qualitative results from the prediction maps and the inference results on the labeled training set. Finally, we investigated how discrepancies in the ‘pgc clover’ predictions is affected by the presence of quadrats in the image using Grad-CAM visualizations and concluded that our dataset would benefit from a more diverse representation of these classes. The results from the baseline models to the SSL FixMatch modeling go to show how powerful it can be to leverage large open source datasets to train accurate image models, all while minimizing the time and monetary costs associated with image annotation.

Furthermore, we deployed the two main DL models on AWS Sagemaker, and then we started development of a simple web application that allows users to login, upload batches of images, and manage image inference jobs for their account. While the application is still under current development, we hope to have it completed by the Fall of 2024 so that users will be able to call the AWS Sagemaker endpoints on their images directly from within the application. To our knowledge, the ‘RegenPGC View’ image analysis pipeline represents one of the first end-to-end deep learning pipelines for cover crop and vegetation segmentation and classification within row cropping plots. While the models and pipeline architecture are still in the nascent stages and could benefit from iterative improvement, this pipeline can be immediately applied to current questions that our research group and other cover-cropping research groups want to answer. We anticipate that the RegenPGC View pipeline and tool will be of great value to researchers who conduct PGC trials, or have projects that overlap with this area of agricultural research.

## 5. Authorship and Acknowledgements

### 5.1. CRediT Attributions

Contributing authors: Bryce Meyering - BM; Brandon Schlautman - BS

- Conceptualization: BM and BS
- Data Curation: BM
- Formal Analysis: BM
- Funding Acquisition: BS
- Investigation: BM
- Methodology: BM
- Project Administration: BM
- Resources: BS
- Software: BM
- Supervision: BS
- Validation: BM
- Writing - original draft: BM
- Writing - review and editing: BM

### 5.2. Funding

Funding for this work was provided by Agriculture and Food Research Initiative Competitive Grant No. 2021-68012-35923 from the USDA National Institute of Food and Agriculture.

## 5.3. Acknowledgements

This work would not be possible without the guidance of Dr. Raj Raman (RegenPGC Project Director, Iowa State University), Dr. Erin Haramoto (Project Co-PD, University of Kentucky), and Dr. Sara Lira (Project Commercialization, Corteva Agriscience). In addition I would like to thank Hallie Sandeen and Kiera Searcy (University of Kentucky); Prathyusha Cheguri and Memis Bilgici (Iowa State University) for all their hard work and efforts acquring the images that comprise the RegenPGC Dataset.

## 5.4. Conflict of Interest Statement

The authors declare they have no conflict of interest in conducting this research.

## 5.5. Data Availability Statement

All data and software used and developed in this project will be made publicly available through a combination of GitHub repositories, Zenodo datasets, and or publicly available web applications at the time of publishing.

## 6. Supplementary Material

### 6.1. Datasheets for Datasets

This section outline the dataset provenance and important considerations for data access[21].

#### 6.1.1 Motivation

- **For what purpose was the dataset created?** This dataset was collected and compiled to support the research of scientists and graduate students on the RegenPGC project in order to estimate the effectiveness of PGC management strategies in PGC establishment, crop yields, and ecological impact.
- **Who created the dataset (e.g**., **which team, research group) and on behalf of which entity (e.g**., **company, institution, organization)?** A wide variety of researchers on the [RegenPGC](https://www.regenpgc.org/our-people/) project including their graduate and undergraduate students were the main drivers of image acquisition and image labeling.
- **Any other comments?** None

#### 6.1.2 Composition

- **What do the instances that comprise the dataset represent (e.g**.,**documents, photos, people, countries)?** Each data instance in the dataset is an image that represents a research field trial plot at a given point in time.
- **How many instances are there in total (of each type, if appropriate)?** For the segmentation model, there are approximately 282 labeled images and 4300 unlabeled images. The dataset used for the object detection model contained XXXXX images, all of which were labeled.
- **Does the dataset contain all possible instances or is it a sample(not necessarily random) of instances from a larger set?** Currently the dataset contains all possible instances that have been collected up to the point of model training.
- **What data does each instance consist of?** The dataset for the segmentation model has labeled and unlabeled instances. Labeled instances contain 1) a.jpg formatted image of size as determined by the device that it was collected on, in addition to a.png formatted binary mask of the same dimensions with unique pixel values representing each of the categorical classes. The object detection dataset also has.jpg images and a csv file with bounding box coordinates for each of the bounding boxes in every image.
- **Is there a label or target associated with each instance?** The segmentation dataset has a labeled and an unlabeled portion while the object detection dataset is fully labeled.
- **Is any information missing from individual instances?** No information is missing besides the absence of labels for the unlabeled data.
- **Are relationships between individual instances made explicit(e.g**., **users’ movie ratings, social network links)?** Not applicable.
- **Are there recommended data splits (e.g**., **training, development/validation**,**testing)?** Yes, these are contained in the data directories.
- **Are there any errors, sources of noise, or redundancies in the dataset?** As with any dataset, there are some minor errors in the labels but none that affect the overall integrity of the labeled data.
- **Is the dataset self-contained, or does it link to or otherwise rely on external resources (e.g**., **websites, tweets, other datasets)?** This dataset was combined with the labeled and unlabeled portions of the GrassClover dataset [**?**] as well as fully labeled CropAndWeed dataset [**?**].
- **Does the dataset contain data that might be considered confidential (e.g**., **data that is protected by legal privilege or by doctor–patient confidentiality, data that includes the content of individuals’ non-public communications)?** No, all of the data that we used is either an internally collected imageset that will be made open source, or an already open source dataset that is available for public use.
- **Does the dataset contain data that, if viewed directly, might be offensive, insulting, threatening, or might otherwise cause anxiety?** No.

#### 6.1.3 Collection Process

- **How was the data associated with each instance acquired?** The images for the RegenPGC dataset, both for the object detection model as well as the segmentation model were collected by RegenPGC affiliated researchers in their research plots and trials.
- **What mechanisms or procedures were used to collect the data(e.g**., **hardware apparatuses or sensors, manual human curation**,**software programs, software APIs)?** Our data was initially collected using standard smartphone cameras (Google Pixel 3a). The majority of the images were captured using the open-source plant phenotyping application, Gridscore, which tags each image with the trial, plot, and treatment information.
- **If the dataset is a sample from a larger set, what was the sampling strategy (e.g**., **deterministic, probabilistic with specific sampling probabilities)?** The labeled images were sampled randomly from the overall collection of images.
- **Who was involved in the data collection process (e.g**., **students**,**crowdworkers, contractors) and how were they compensated (e.g**.,**how much were crowdworkers paid)?** Graduate students, public university employees, non-profit researchers, and industry professionals all contributed to collecting images and the labeling process.
- **Over what timeframe was the data collected?** The majority of the images were collected in the time of field planting to harvest over a period of two growing seasons.
- **Were any ethical review processes conducted (e.g**., **by an institutional review board)?** No review process was conducted for this data collection.

#### 6.1.4 Preprocessing/cleaning/labeling

- **Was any preprocessing/cleaning/labeling of the data done (e.g**., **discretization or bucketing, tokenization, part-of-speech tagging, SIFT feature extraction, removal of instances, processing of missing values)?** Images that were not suitable for modeling, i.e. images that were taken at a highly oblique angle or were too blurry for modeling were removed from the dataset. Some preprocessing such as normalization and resizing were applied before modeling, but were performed on the fly and not stored. The RegenPGC image masks needed to be converted to a common set of labels that are compatible with the masks in the CropAndWeed and GrassClover datasets. For instance, the CropAndWeed labels included many species level weed labels whereas our model only included a ‘broadleaf-weed’ category. This necessitated the weed species subcategories to be aggregated up into a larger supercategory.
- **Was the “raw” data saved in addition to the preprocessed/cleaned/labeled data (e.g**., **to support unanticipated future uses)?** Yes the raw images and labels were all stored in addition to the processed images.
- **Is the software that was used to preprocess/clean/label the data available?** All of the labels that we generated for the RegenPGC dataset were generated using the commercial software, Labelbox. Preprocessing of the images was done in Python using OpenCV and Albumentations.
- **Any other comments?** None.

#### 6.1.5 Uses

- **Has the dataset been used for any tasks already?** No, this is the first use of these images.
- **Is there a repository that links to any or all papers or systems that use the dataset?** No, this is the first use of this dataset. All of the code for the models and the application that use this data are available.
- **What (other) tasks could the dataset be used for?** The images could be relabeled and used for segmenting specific weed species.
- **Is there anything about the composition of the dataset or the way it was collected and preprocessed/cleaned/labeled that might impact future uses?** The data is very specific to certain species of perennial grasses and clovers and modeling this data may not be applicable to other species of grasses.
- **Are there tasks for which the dataset should not be used?** None.
- **Any other comments?** None.

#### 6.1.6 Distribution

- **Will the dataset be distributed to third parties outside of the entity (e.g**., **company, institution, organization) on behalf of which the dataset was created?** The dataset will be made publicly available for download.
- **How will the dataset will be distributed (e.g**., **tarball on website, API, GitHub)?** We plan to host all of the images and labels in a dedicated Zenodo repository. Zenodo releases a concept DOI that always points to the newest version of the dataset as well as version specific DOIs.
- **When will the dataset be distributed?** At the time of publishing.
- **Will the dataset be distributed under a copyright or other intellectual property (IP) license, and/or under applicable terms of use(ToU)?** This will be distributed under some creative commons terms of use so that it is available for use.
- **Have any third parties imposed IP-based or other restrictions on the data associated with the instances?** None.
- **Do any export controls or other regulatory restrictions apply to the dataset or to individual instances?** None.
- **Any other comments?** None.

#### 6.1.7 Maintenance

- **Who will be supporting/hosting/maintaining the dataset?** The Land Institute, a 501(c)(3) non-profit agricultural research organization will take on the responsibility of maintaining and updating the dataset.
- **How can the owner/curator/manager of the dataset be contacted(e.g**., **email address)?** The current curator of the dataset should be contacted by email.
- **Is there an erratum?** None is currently available.
- **Will the dataset be updated (e.g**., **to correct labeling errors, add new instances, delete instances)?** We plan to update the data as we label more images in the dataset. The Zenodo repository containing the dataset will be appropriately versioned as the dataset is updated.
- **If the dataset relates to people, are there applicable limits on the retention of the data associated with the instances (e.g**., **were the individuals in question told that their data would be retained fora fixed period of time and then deleted)?** None of the images or labels in the dataset pertain to people.
- **Will older versions of the dataset continue to be supported/hosted/maintained?** Since Zenodo repositories can be versioned and stored, all of the previous versions of the data will be available as long as Zenodo can provide support.
- **If others want to extend/augment/build on/contribute to the dataset, is there a mechanism for them to do so?** They can contact the dataset curators by email or reach out to curator team for the Zenodo repository to discuss how to contribute to the furthering of the dataset.
- **Any other comments?** None.

## Footnotes

1 1909, U.S. Bureau of Soils

2 RegenPGC is supported by Agriculture and Food Research Initiative Competitive Grant No. 2021-68012-35923 from the USDA National Institute of Food and Agriculture.

## Notes

### Competing Interest Statement

The authors have declared no competing interest.

